# sgcocaller and comapr: personalised haplotype assembly and comparative crossover map analysis using single-gamete sequencing data

**DOI:** 10.1101/2022.02.10.479822

**Authors:** Ruqian Lyu, Vanessa Tsui, Wayne Crismani, Ruijie Liu, Heejung Shim, Davis J. McCarthy

**Affiliations:** Bioinformatics and Cellular Genomics, St. Vincent’s Institute of Medical Research,9 Princes Street, Fitzroy Victoria, 3065, Australia; Melbourne Integrative Genomics/School of Mathematics and Statistics, Faculty of Science, The University of Melbourne, Building 184, Royal Parade, Parkville, Victoria,3010, Australia; DNA Repair and Recombination Laboratory, St Vincent’s Institute of Medical Research,9 Princes Street, Fitzroy Victoria, 3065 Australia; The Faculty of Medicine, Dentistry and Health Science, The University of Melbourne, Victoria 3010 Australia

**Author notes:** H.S. and D.J.M. supervised this work.

## Abstract

Profiling gametes of an individual enables the construction of personalised haplotypes and meiotic crossover landscapes, now achievable at larger scale than ever through the availability of high-throughput single-cell sequencing technologies. However, high-throughput single-gamete data commonly have low depth of coverage per gamete, which challenges existing gamete-based haplotype phasing methods. In addition, haplotyping a large number of single gametes from high-throughput singlecell DNA sequencing data and constructing meiotic crossover profiles using existing methods requires intensive processing. Here, we introduce efficient software tools for the essential tasks of generating personalised haplotypes and calling crossovers in gametes from single-gamete DNA sequencing data (sgcocaller), and constructing, visualising, and comparing individualised crossover landscapes from single gametes (comapr). With additional data pre-possessing, the tools can also be applied to bulk-sequenced samples. We demonstrate that sgcocaller is able to generate impeccable phasing results for high-coverage datasets, on which it is more accurate and stable than existing methods, and also performs well on low-coverage single-gamete sequencing datasets for which current methods fail. Our tools achieve highly accurate results with user-friendly installation, comprehensive documentation, efficient computation times and minimal memory usage.

## Introduction

Meiosis is a process required during sexual reproduction that generates gametes—egg or sperm cells—that contain half of the chromosomes of the parent cell (1). Chromosome segregation and meiotic crossovers create abundant genetic diversity in the offspring. Meiotic crossovers are also required for accurate chromosome segregation (1), and reduced crossover rates are linked to increased risk for trisomy 21 (2, 3) and infertility (4, 5). Past studies have shown that meiotic crossover rates and distributions vary greatly among species, sexes and even individuals (6–11). Variation in genetic factors, such as *PRDM9*, changes the distribution of crossover hotspots in human and mouse (6, 8, 12, 13). Crossover-regulating genes also limit overall crossover rates (14–16). Populations of different demographic backgrounds show differences in recombination landscapes, for example hotspot locations differ in African-American genetic maps compared with those from Europeans and West Africans (17, 18). It has also been shown that meiotic crossovers vary among sexes and crossover hotspots can be sex-specific (7, 10, 19).

Single-cell DNA sequencing of gametes collected from an individual can be used to construct personalised meiotic crossover distributions (9, 11). The scalability of modern single-cell assays, especially droplet-based platforms, has made it possible to profile thousands of sperm cells per individual in one experiment (9). Assaying gametes from females is substantially more challenging at scale, but nevertheless technological developments will make large-scale single-gamete sequencing more common across sexes and organisms. The standard preprocessing pipeline of a single-cell DNA sequencing dataset generates a BAM (BinaryAlignment/Map) file that contains the mapped and cell-barcoded DNA reads from all single cells. Haplotyping these singlecell genomes with existing tools requires the steps of demultiplexing the single-cell reads according to their cell barcodes into thousands of intermediate files before applying haplotyping methods (20). While processing can be parallelised across cells, there are many opportunities to improve upon the use of methods developed for bulk DNA-seq data when analysing single-cell DNA-seq data by avoiding multiple reading and writing of millions of sequence reads. Even more importantly, there are substantial opportunities to improve on the accuracy of existing methods for haplotype reconstruction and genetic map construction from single-gamete data. Single-gamete DNA sequencing data usually have low depth of coverage across the genome per gamete, especially when generated using high-throughput droplet-based single-cell sequencing protocols. However with a group of gametes sequenced, the single nucleotide polymorphism (SNP) linkage in the gametes offers enough information for constructing chromosome-level phased haplotypes of the individual.

Therefore, we here introduce sgcocaller, an efficient command-line toolkit to directly process the large singlecell DNA-sequencing alignment files produced by the current standard pre-processing pipelines for personalised haplotype construction and single-gamete crossover identification. sgcocaller is also applicable to individual gametes sequenced by bulk DNA sequencing methods with minimal dataset preprocessing. We also introduce an associated Bioconductor/R package, comapr, for the construction, visualisation, and statistical analysis of crossover landscapes. Working in the Bioconductor ecosystem enables comapr to integrate seamlessly with other commonly used packages specialised in analysing biological datasets. Our new tools offer substantial improvements in accuracy of haplotype reconstruction from single-gamete sequencing data as well as greatly enhanced computational efficiency. With these tools, an easy-to-apply and endto-end workflow for personalised haplotype assembly and comparative crossover map analysis has been made possible.

## Materials and Methods

### Overview of sgcocaller and comapr

We have implemented a toolset for processing large-scale single-cell DNA sequence data from gametes with modules to construct the personalised haplotypes of the individual and to call crossovers for individual gametes. The toolset consists of two components, sgcocaller and comapr (see), for complementary tasks. These packages have been implemented using appropriate programming languages (Nim and R) and released in multiple release formats to suit the expected use-case scenarios (Fig. 1).

**Fig. 1.**
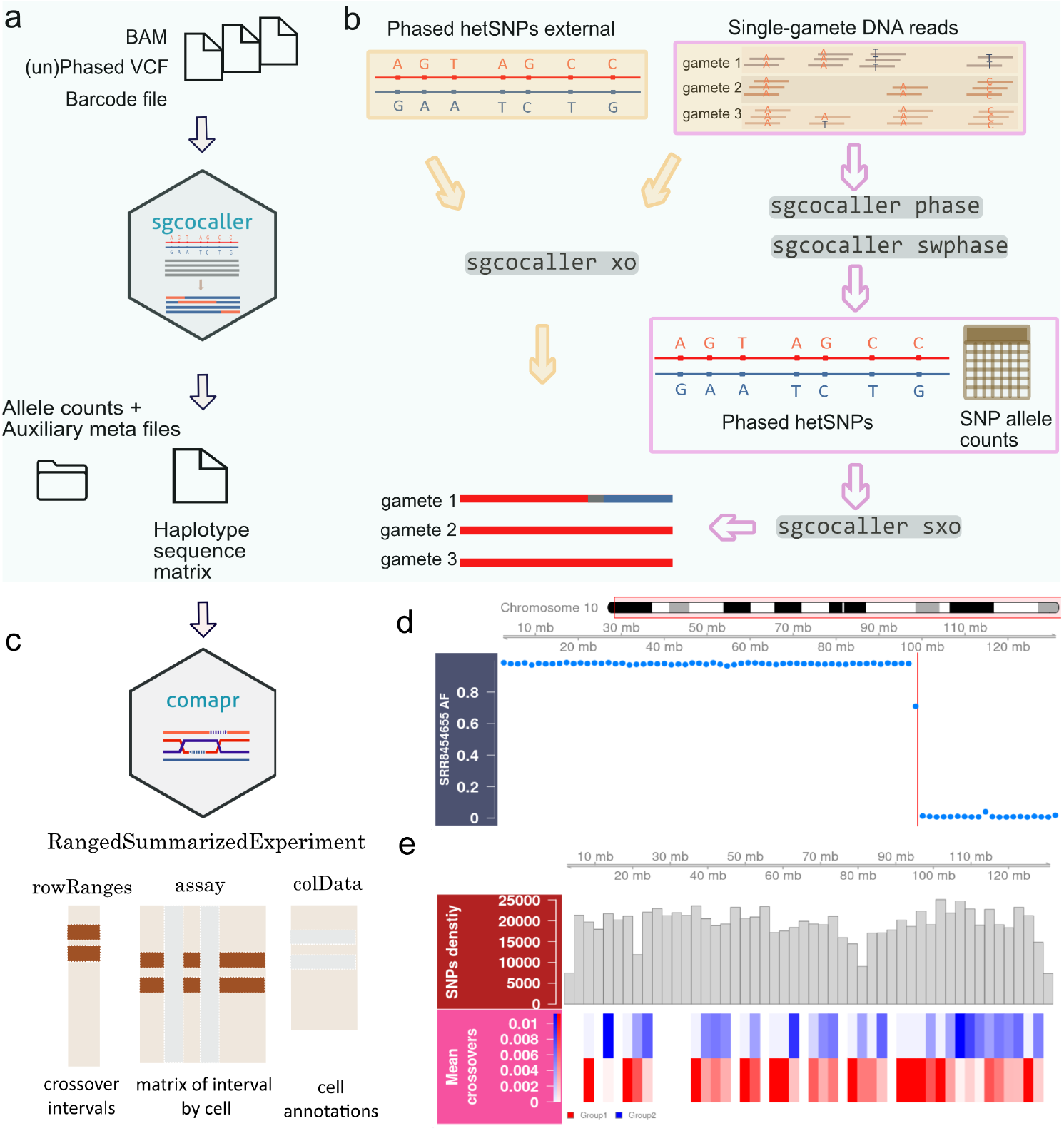
Overview of sgcocaller and comapr workflow including file and data flows of sgcocaller and comapr, as well as example plots generated by comapr. **a)** sgcocaller takes the aligned DNA reads from all gametes, the list of hetSNPs (phased or unphased) and the cell barcode list and produces the single-gamete haplotype sequence in sparse matrix. **b)** The diagram shows the main function of sgcocaller which is to resolve single gametes’ haplotypes from DNA reads using phased hetSNPs obtained from external sources (left) or from applying *sgcocaller phase* (right) by the two demonstrated workflows. **c)** Output files from sgcocaller can be further processed and analysed by comapr, which adopts the *RangedSummarizedExperiment* as the main data structure to enable seamless integration with existing Bioconductor packages. **d)** Alternative allele frequencies plot for binned windows with crossover regions highlighted (vertical bar) from one selected gamete cell. **e)** The heatmap plot of crossover counts for two groups of mouse sperm cells with SNP density plot in histogram on top (both plots used the mouse sperm dataset (21)).

### sgcocaller: single-gamete crossover caller

To suit the requirements of processing large alignment files containing DNA sequence read data for hundreds or thousands of individual gametes, sgcocaller has been designed as a commandline tool (CLT) implemented using the programming language Nim (https://nim-lang.org/). The software imports the *hts-nim* library (22) for fast processing of large DNA alignment files and variant calling files. It also imports the Rmath library (in programming language C from the R-project (23)) to use the well-defined distribution functions. To interface with the C-based library, a wrapper package (in Nim) was created and implemented as the *distributions* package that is openly accessible (see). In crossover-calling scenarios with known phased haplotypes such as F1 hybrid samples generated by crossing known reference strains of species, and cases in which the haplotypes of donors are provided in an input VCF file, *sgcocaller xo*—the crossover calling module—can be applied directly. It requires three input files to call crossovers: the mapped BAM file with cell-barcoded DNA reads from all gametes, the VCF file of phased heterozygous SNP (hetSNP) markers and the list of cell barcodes (Fig. 1a). When the phased haplotype information is not available from external sources, sgcocaller also offers a phasing module, *sgcocaller phase*, that is based on the SNP linkage data inherent in read data from the individual gametes to produce the personalised whole-chromosome haplotypes. To call crossovers using outputs of *sgcocaller phase, sgcocaller sxo* is recommended as it uses the intermediate files generated by *sgcocaller phase* including the phased haplotypes and the allele specific read count matrices as inputs, which avoids double handling of BAM/VCF files (Fig. 1b). Our tool is engineered to operate directly on the output of common single-cell data processing pipelines such as CellRanger, STARSolo (24) and similar. With minimal preprocessing, sgcocaller can also work with bulk-sequencing samples as demonstrated in the application example with the mouse single-sperm dataset (see).

### comapr: crossover analysis for construction of genetic distance maps in R

The second software component, comapr, serves as a post-processing tool that includes functions for finding crossover positions, quantifying crossover rates across groups, and conducting comparative analyses after the sequences of haplotypes are inferred by *sgcocaller xo* (or *sgcocaller sxo*) for each chromosome in gametes. It is implemented as an open-source Bioconductor/R package, which offers easy integration with other packages in Bioconductor for analysis of biological data and statistical testing. comapr includes functions to directly parse output files generated by *sgcocaller xo* or *sgcocaller sxo* with tunable parameters to systematically filter out potential false positive crossover calls and create structured data objects to represent the crossover information across all cells (Fig. 1c). comapr provides quality checking functions and visualisations to understand the features of the underlying dataset and for choosing sensible filtering thresholds. It enables convenient plotting of summary plots such as number of crossovers per sample group, converting crossover rates to genetic distances in units of centiMorgans, and plotting genetic distances over chromosomes or the whole genome (see). comapr integrates with the genomic visualisation package Gviz (25) and easily generates alternative allele frequency plots with crossover intervals highlighted or with genomic feature tracks overlaid on top of identified crossover tracks (Fig. 1d,e). To facilitate easy statistical significance testing for comparisons of crossover rates in gamete groups, such as gametes collected from individuals with different genetic backgrounds or from different experimental groups, comapr implements two resampling based methods, bootstrapping and permutation testing.

### sgcocaller phase and sgcocaller swphase

Detecting crossovers in gametes’ genomes requires the haplotypes of the individual’s genome. When the individual’s haplotypes are not available from other sources, sgcocaller offers a module *sgcocaller phase* that is able to produce the phased hetSNPs from unphased hetSNPs using the available single-gamete data (Fig. 1b). *sgcocaller phase* uses SNP markers’ co-appearance information across all gametes to generate the chromosome-scale haplotype of the individual from single-gamete data, an idea that has also been applied in a previous study (26) (Fig. 2a). To increase the algorithm’s efficiency and fully utilize the known biological mechanisms of meiosis and meiotic crossovers, the phasing algorithm in *sgcocaller phase* first finds a template cell (a cell putatively without crossovers) and uses the cell’s genotype sequence as the template haplotype (Fig. 2a; see Supplementary Methods). It next fills in the missing SNPs in the template haplotype using SNP linkage information from other gametes to increase the completeness of the phased haplotypes. The inference of missing SNPs is based on the fact that meiotic crossovers are low frequency events across chromosomes and crossover positions are sparse. The SNP linkages in small chromosome regions across all haploid gametes are therefore reliable for reconstructing the donor’s haplotypes (see Supplementary Methods).

**Fig. 2.**
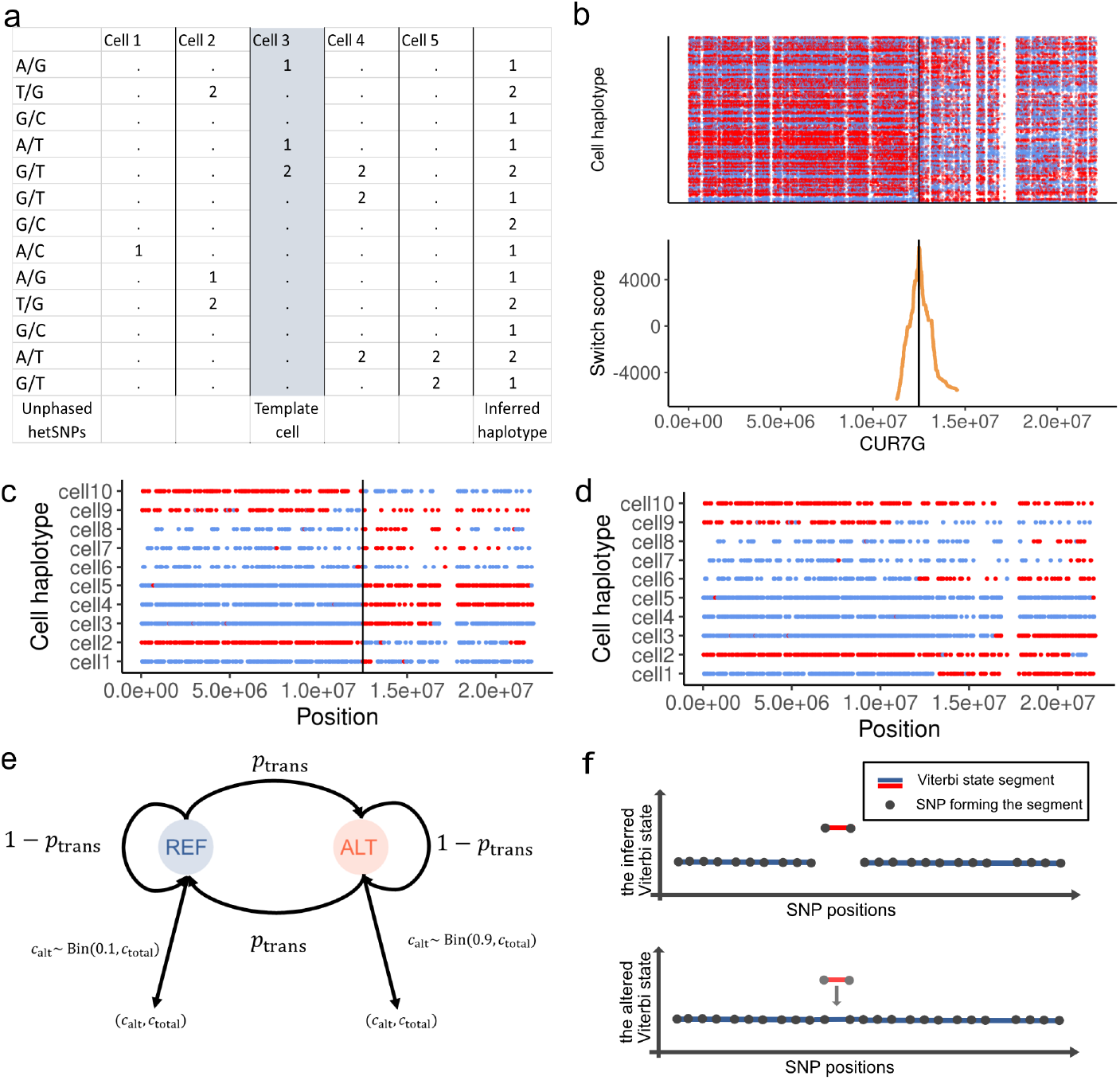
*sgcocaller phase* and *sgcocaller swphase* generate personalised haplotypes from single-gamete data **a)** Genotype matrix of an example of five gametes regarding the list of the hetSNPs with the chosen template cell highlighted. Template cell’s genotype sequence is used as the backbone for generating the inferred personal haplotype. **b)** (Top) Cell genotypes along the list of SNPs were plotted and colored by which haplotypes they were matched with (the inferred template haplotype or its complimentary haplotype). (Bottom) Switch scores were calculated and plotted for SNPs in the inferred haplotype sequence, which were based on the genotypes of the flanking SNPs across all cells. The switch error position was found as the identified peak in switch scores (the black line). **c**,**d)** Genotype sequences of ten cells were plotted and colored by whether the SNP’s genotype matching the inferred template haplotype or its complimentary haplotype for the case when there was a switch error in the inferred haplotype with the error position indicated by the black vertical line (c), and when there was no switch errors (d). Plots in b,c,d were generated using the apricot gametes dataset (20). **e)** Diagram of the two-state (REF and ALT) hidden Markov model implemented in *sgcocaller xo*. The two possible alleles at each hetSNP site can be referred as REF or ALT arbitrarily. Binomial distributions have been adopted for modelling the relationship of observed allele counts and the underlying haplotype states of hetSNPs. The transition probability between two states (*p*_trans_, the distribution parameters (e.g., 0.1, 0.9) are configurable options when running *sgcocaller xo* (see). **f)** Diagram of the Viterbi state segment with the inferred hidden states (output from *sgcocaller xo*) and the altered hidden states (artificially generated by reversing the inferred states of the SNPs in the underlying segment). Segments are colored by their states. Dots represent the SNPs forming the segments.

*sgcocaller phase* also generates auxiliary files for conveniently plotting diagnostic plots that help to inspect the quality of the inferred haplotypes and check for switch errors (Fig. 2c). Cells in the context of this manuscript are haploid gametes unless otherwise specified.

Since crossovers are low frequency events at each site across the gametes, switch errors in the inferred haplotypes can be identified by inspecting the diagnostic plot. When the plotted genotype sequences of all cells show existence of crossovers at the same position, it indicates a “crossover” or switch error has occurred in the inferred haplotype (Fig. 2c,d). Therefore, we have included a switch error correction module *sgcocaller swphase* for finding and correcting the switch errors if any in the phased haplotype constructed by *sgcocaller phase*. It produces switch scores for a selected list of SNPs with a higher risk of having switch errors and identifies the switch positions to generate the corrected haplotype (Fig. 2b; see Supplementary methods).

### sgcocaller xo and sgcocaller sxo

The crossover calling module *sgcocaller xo* infers the haplotype states in the haploid genomes of gametes. Two haplotypes (represented by the list of red or blue alleles on each chromosome in Fig. 1b) can be found in the diploid genomes of individuals and the gametes from the individual inherit a combination of the two haplotypes (Fig. 1b). Gametes have haploid genomes, therefore theoretically there is only one type of allele that can be observed at each SNP position in each gamete. However, due to technical noise such as mapping artefacts, there can be a substantial number of SNP sites with two types of alleles found but with biased ratio towards the true allele type in the genome. To suppress the potential noise in the dataset, *sgcocaller xo* implements a two-state hidden Markov model (HMM) and adopts binomial distributions for modelling the emission probabilities of the observed allele read counts (Fig. 2e; see Supplementary Methods).

With this HMM, the commonly used dynamic programming method, the Viterbi algorithm (27), is implemented in *sgcocaller xo* to solve for the most probable hidden state sequence for each chromosome in each cell. Since the two hidden states in the HMM represent the haplotype origins of DNA segments (represented by allele types of a list of SNPs) in the gametes’ genomes, the transition from one state to another in the hidden state sequence between two SNP markers corresponds to a crossover detected. As currently designed, sgcocaller is intended to work on data of gametes collected from diploid individuals.

The *sgcocaller sxo* module supplements the crossover calling module *sgcocaller xo*. It runs the same core function as *sgcocaller xo* but uses the generated outputs (i.e., allele count matrices and the phased haplotypes) from *sgcocaller phase* instead of the original BAM and VCF files, thus eliminating unnecessary double handling of large DNA sequencing data. In addition to applying the HMM to the allele counts and using the Viterbi algorithm for inferring the hidden state sequences, we also included calculation of a crossover confidence score, which is the log-likelihood ratio of the Viterbi state segment. The log-likelihood ratio of a segment is derived by finding the difference between the log-likelihood of the segment given the inferred state and the log-likelihood of the segment given the altered state (log-likelihood of the first path given the data minus the log-likelihood of the second path; Fig. 2f; see Supplementary Methods).

### Scope of sgcocaller

Although sgcocaller has been optimised to work on barcoded large-scale single-cell DNA sequencing data with outputs in formats of sparse matrices, it can also—with simple preparation—be applied to sequencing reads from one cell or bulk DNA sequencing samples for which the cell barcodes are not available.

For single-gamete DNA sequencing datasets generated using bulk-like protocols (individual sequencing libraries that generated separate sets of reads for gametes), sgcocaller can still be applied, and we have released open-access tools for preparing such data to make it compatible with the *sgcocaller-comapr* pipeline. We provide examples of applying sgcocaller on such datasets (21, 28) (see).

### comapr: implementation

#### Data structure

The ‘RangedSummarizedExperiment’ class defined in the Bioconductor R package ‘SummarizedExperiment’ (29) is used as the main data structure in comapr for storing the SNP interval regions and the number of called crossovers per cell (Fig. 1). The data slot ‘rowRanges’ is used for storing the SNP intervals, whereas the cell-level information is stored in the ‘colData’ slot. Using ‘RangedSummarizedExperiment’ also enables comapr to conveniently integrate with the various genomic coordinate plotting functions in the Gviz package (30).

#### Resampling-based functions for testing differences in crossover profiles

To test for differences in the number of crossovers between any two groups of cells, we have implemented re-sampling methods in comapr, specifically permutation and bootstrapping tests (31–33). The two resampling-based functions in comapr are able to either calculate empirical p-values (permutation testing) or bootstrap confidence intervals for the estimate of the group differences (see Supplementary Methods).

### Preprocessing public datasets

#### Mouse sperm dataset

Raw fastq files of 217 mouse sperm cells were available from accession GSE125326 in the Gene Expression Omnibus (21), and fastq files of 194 mouse sperm cells were downloaded. The downloading process of the rest 23 cells failed and they were not analysed. We applied `fastp-v0.20.1` (34) on the raw reads for filtering out low quality reads and adapter trimming, before applying `minimap2-2.7_x64-linux` (35) for aligning the reads to the mouse reference genome mm10. The mapped reads were further processed by `GATK MarkDuplicates` and `GATK AddOrReplaceReadGroup` from the `GATK-v4.2` pipeline (36). Read sorting and indexing were performed using `samtools-v1.10` (37, 38). A customised open-access tool `appendCB` (see) was applied to add barcode sequences to each sperm’s DNA reads with tag CB using its SRR sequence. The DNA reads in each sperm sample were sub-sampled to retain half of the reads and merged into one barcode-tagged large BAM file that was analysed in this study as the mouse sperm dataset. The merging and indexing of BAM files were achieved using `samtools-v1.10` (37). We followed the main steps described in the original study for finding heterozygous SNPs in the mouse donor’s genome (21). We first called variants *de novo* on the bulk sperm sample ‘SRR8454653’ (sequenced DNA reads of pooled multiple sperm) using `GATK-HaplotypeCaller`. Only the hetSNPs with mapping quality score larger than 50, and depth of coverage within the range of 10 to 80 were kept (MQ>50 AND DP>10 AND DP<80).

Since the mouse donor was an F1 hybrid (C57BL/6J X CAST/EiJ), the list of reference hetSNPs was downloaded (`CAST_EiJ.mgp.v5.snps.dbSNP142.vcf.gz` and `C57BL_6NJ.mgp.v5.snps.dbSNP142.vcf.gz`) from the Mouse Genome Project (39). The called variants in sample SRR8454653 were further filtered to only keep the positions which were called as homozygous alternative (`GT==1/1`) in `CAST_EiJ.mgp.v5.snps.dbSNP142.vcf.gz` and not overlapping with variants in `C57BL_6NJ.mgp.v5.snps.dbSNP142.vcf.gz`. Scripts are publicly available (see).

#### Further sub-sampled mouse sperm dataset

To generate a mouse sperm dataset that mimics the coverage level of a typical single-gamete DNA sequencing dataset (e.g., apricot gamete dataset), the merged mouse sperm BAM file (as described before in section) was sub-sampled using `samtools-v1.10` (37) to a fraction of 0.15 to yield the further sub-sampled mouse sperm dataset.

#### 10X scCNV apricot gametes

Two experiments were conducted to sequence the apricot gametes in the original published study (20). The pre-aligned BAM files (of two experiments) were downloaded from European Nucleotide Archive (ENA) under accession number “PR-JEB37669”. The downloaded pre-aligned BAM files were converted to fastq reads with `samtools-v1.10`. To keep the cell barcode information for each DNA read in the converted fastq files, the cell barcode sequence was appended to each fastq read’s sequence name. Reads in fastq files were then mapped to the published haploid genome “Currot” using `minimap2-2.7_x64-linux`. The identification of hetSNP markers was performed by running `bcftools-v1.10` on the pooled DNA reads from the two experiments. The identified hetSNPs were filtered using the same command as from the original study(20) (`QUAL > 200 & FORMAT/DP < 280 & FORMAT/DP > 120 & FORMAT/GT==‘0/1’ & (FORMAT/AD[0:1])/(FORMAT/DP) > 0.38 & (FORMAT/AD[0:1])/(FORMAT/DP) < 0.62`). Gametes from the two experiments were merged in the analysis and gametes with barcode collisions were removed. Scripts are publicly available (see).

### Phasing performance comparison

Performance of *sgcocaller phase*|*swphase* was compared to Hapi (40) on the human sperm dataset (28),the apricot gametes generated by the 10X scCNV protocol (20), the mouse sperm dataset (21) and the further sub-sampled mouse sperm dataset (see Results:).

#### Human sperm cell dataset

We constructed 11 datasets by leaving one sperm out from the 11 available sperm cells in the human sperm dataset (28). The list of unphased hetSNPs and phased results (used as ground truth) were downloaded from the supplementary dataset shared in the published study (28). The parameter settings of each method were kept consistent when applying the two methods on the 11 datasets. Hapi was run following the tutorial example and the allowNA parameter in imputing missing genotypes function was set to 3 (see Supplementary Methods).

#### 10X scCNV apricot gametes

For the apricot dataset (20), 10 datasets were constructed by leaving out a different 10% of the gametes each time. The haploid genome assembly “Currot” published previously was used as the haplotype ground truth. The alleles in the list of called hetSNPs was swapped for every other position to create the list of unphased hetSNPs. Due to the sparsity of the dataset, the parameters were adjusted for both methods (see Supplementary Methods).

#### Mouse sperm dataset

For the mouse sperm dataset (21), 10 datasets were constructed by leaving out a different 10% of the gametes each time. The phase of the known mouse strains were used as the ground truth. The alleles in the list of called hetSNPs was swapped for every other position to create the list of unphased hetSNPs (see Supplementary methods). Different input options were applied when running the two methods on the further sub-sampled mouse sperm dataset (see) to account for the different coverage profiles of the datasets (see Supplementary Methods).

#### Calling crossovers

We applied sgcocaller and comapr on public datasets to demonstrate the application of the software tools (see). The application examples cover different use-case scenarios: 1) a mouse sperm dataset (21) for which the haplotypes of the donors were known and 2) the apricot gamete dataset with haplotypes of the donor inferred by *sgcocaller phase* before calling crossovers using *sgcocaller sxo*. For the apricot dataset, crossover results when using the phased hetSNPs obtained from the published study (20) were also generated using *sgcocaller xo* to compare the differences in crossover calling results between using the haplotypes inferred by *sgcocaller phase* and the known haplotypes from the original study (see Supplementary Methods).

The output files from *sgcocaller xo* and *sgcocaller sxo* (see Supplementary Methods) were further processed using functions from comapr for cell filtering, false positive crossover filtering and genetic distances calculation and visualisation (see Supplementary Methods and public code repositories).

## Results

### sgcocaller generates highly accurate phasing results

We applied *sgcocaller phase* (and *sgcocaller swphase* when needed) on an apricot pollen dataset generated with the 10X scCNV protocol (a droplet-based single-cell DNA sequencing protocol) (20). *sgcocaller swphase* can be run selectively and only for cases when switch errors are identified in the inferred haplotype by *sgcocaller phase*. We refer to the final phasing results as generated by *sgcocaller phase* although *sgcocaller swphase* may have been applied. The phasing results of *sgcocaller phase* demonstrate very high concordance with the haplotypes from the published assembly from the same study (Fig. 3a,b). To evaluate performance of the phasing module in sgcocaller, the phasing accuracy was calculated using the fraction of concordant hetSNPs between the haplotype inferred from *sgcocaller phase* and the published haplotype. The hetSNPs were grouped into bins with 100 consecutive SNPs in each bin. The proportion of SNPs matching the published haplotype in 100-SNP bins concentrated at 1.0 across eight chromosomes indicating high phasing accuracy across the genome (Fig. 3a). When defining the alternative alleles as the alleles with lower total abundance in each bin and summarising the alternative allele reads frequencies (AAF) in 1,000-SNP bins, we found that using the haplotypes generated by *sgcocaller phase*|*sgcocaller swphase* resulted in more bins with AAF valued at zero than using the published haplotype (Fig. 3b,c). These results suggest *sgcocaller phase*|*sgcocaller swhase* generated haplotypes that are more concordant with the single-gamete read data than the published assembly.

**Fig. 3.**
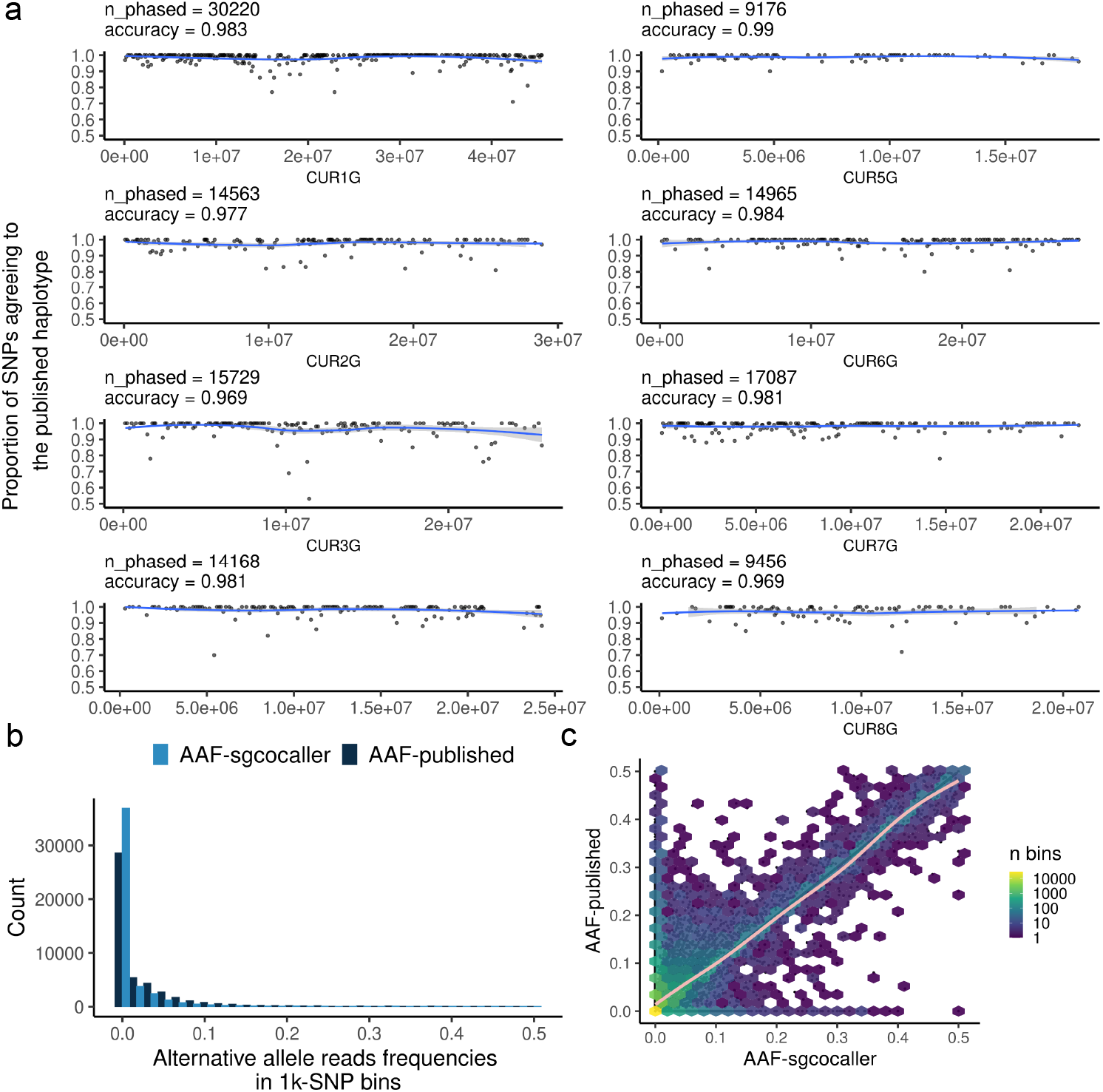
Comparisons between haplotype inferred by *sgcocaller phase* versus the published haplotype using the apricot dataset (20) **a)** Proportion of SNPs agreeing with the published haplotype in 100-SNP bins across eight chromosomes from the using the apricot dataset (20). The number of phased SNPs (*n_phased*) by *sgcocaller phase* and phasing accuracies (*accuracy*) were printed on top of each panel. Smoothed curves were fitted by using “loess” method from R (23, 41). **b)** The histogram of alternative allele reads frequencies (AAF) for bins of 1,000 SNPs were plotted from all chromosomes colored by results using the haplotype generated by *sgcocaller phase* (AAF-sgcocaller) or the published haplotype (AAF-published). More bins with AAF valued at zero when using the haplotype by *sgcocaller phase* suggested that the haplotype from *sgcocaller phase* was more concordant with the single-gamete data. **c)** The two alternative read frequencies (AAF-sgcocaller and AAF-published) for each bin were plotted in scatter plot with AAF-sgcocaller on the x axis and AAF-published on the y axis. Hexagons were plotted overlaying the scatter plot with color scale indicating counts of bins covered by the hexagon area. Fitted curve that aligned with the diagonal line was plotted using the method “gam” from ggplot2 (23, 41, 42). Alternative allele in these plots are always the alleles with fewer read counts in each bin.

### Application of sgcocaller and comapr on public datasets

#### Bulk DNA sequencing dataset of single mouse sperm cells

We applied sgcocaller and comapr to haplotype sperm cells from a published DNA sequencing dataset of individual mouse sperm cells collected from an F1 hybrid mouse (C57BL/6J X CAST/EiJ) (21). The raw sequencing dataset was downloaded from GEO (Gene Expression Omnibus) with accession GSE125326 (see Supplementary Methods) and the workflow with detailed steps for preprocessing the raw reads and the execution of *sgcocaller*|*comapr* is publicly available (see). The crossovers called by *sgcocaller*|*comapr* (with a mean number of crossovers of 12 per sperm across autosomes) were highly consistent with the crossovers called from the original paper (Fig. 4). There were 7 cells (4%) that were called with a different number of crossovers compared to the original paper (Fig. 5a). We also compared the genetic distances calculated when using the estimated crossover rates by *sgcocaller xo* and the published study (21) in 10 Mb intervals, and it showed perfect alignment between the two methods (Fig. 5b and Fig. S1). We demonstrate and provide reproducible code for running the functions to generate plots of the number of crossovers per sperm cell (Fig. 4a), genetic distances in the chosen size of chromosome bins along chromosomes (Fig. 4b), and the number of crossovers (COs) identified per chromosome (Fig. 4c) in the pubic repository (see). To demonstrate the functions in comapr that test for differences in groups of cells, such as cells from different individuals, we divided the sperm cells into two groups randomly. The crossover counts and crossover rates of each group are calculated over the SNP intervals via function `countCOs` and genetic distances are derived by the `calGeneticDist` function that applies a user-selected mapping function such as the Kosambi mapping function (43). Comparative plots can be generated for the two groups including genetic distance plots for binned intervals along the chromosome and the cumulative genetic distances along a chromosome or whole genome (Fig. 4d,e,f). Resampling-based testing functions, bootstrapping and permutation, have been called to assess the group differences statistically (Fig. 4g,h). The bootstrapping testing was conducted via `bootstrapDist` function. The bootstrap 95% confidence interval of the group differences was calculated. The interval included zero, indicating that there was not enough evidence to support a difference in mean crossover numbers between the two groups at a significance level of 0.05, which was expected because the group labels of sperm cells were randomly assigned (Fig. 4g). The permutation testing via `permuteDist` returned the same conclusion with a permutation p-value *>* 0.05 (Fig. 4 and see).

**Fig. 4.**
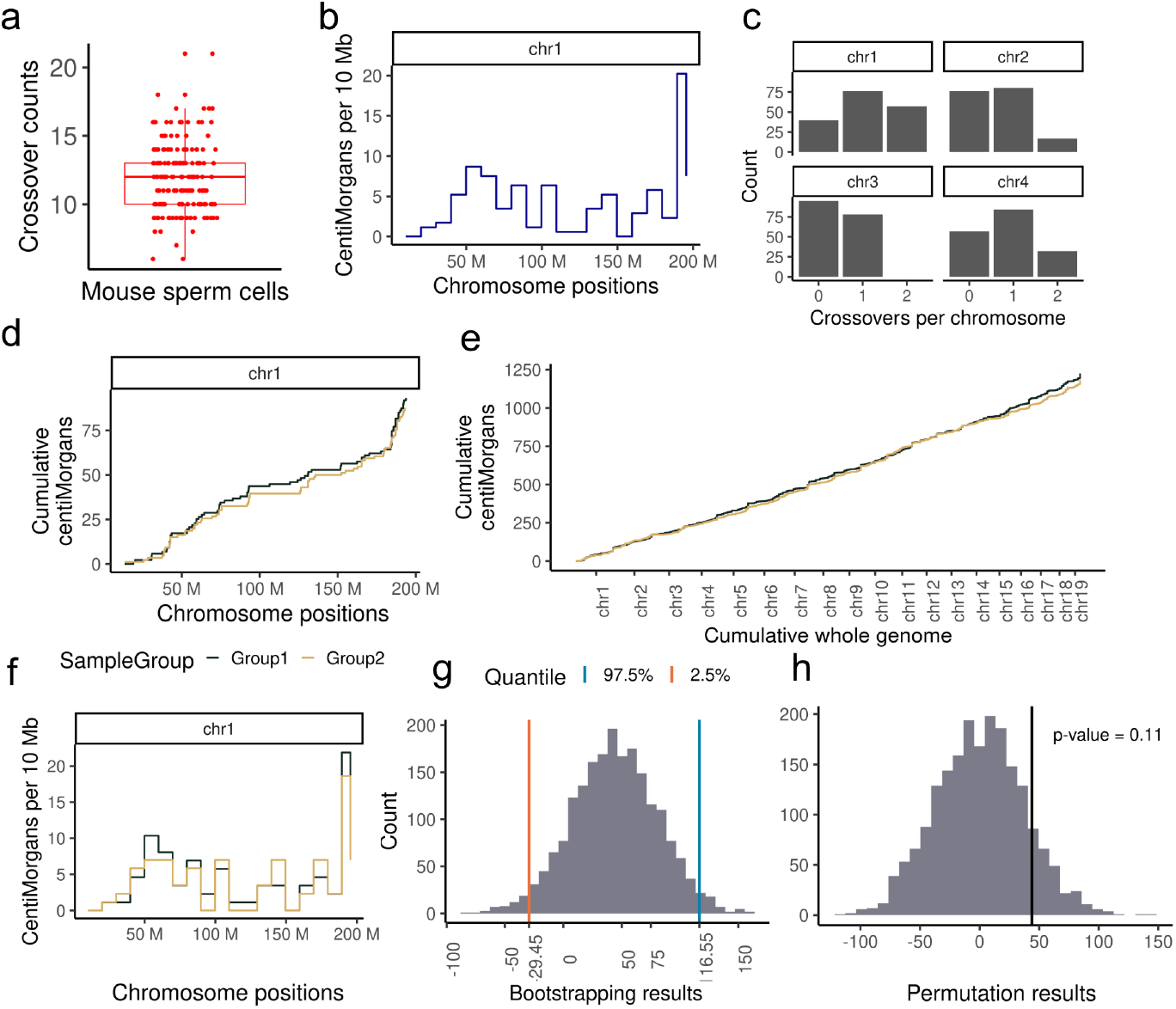
Application of sgcocaller and comapr on mouse sperm dataset (21) comapr enables easy plotting of crossover summary statistics. **a)** The distribution of crossover counts per sperm (n = 173 sperm cells). **b)** The genetic distances (in centiMorgans, which were derived from observed crossover rates by applying the mapping function Kosambi (43) with implemented function in comapr) plotted for every 10 megabase bin along chromosome 1. It has shown a low to zero crossover rate around the centromere region (125M). **c)** The frequency of crossover counts for four chromosome are plotted for all sperm cells. **d**,**e)** The cumulative genetic distances are plotted for chromosome 1 and cumulatively whole genome for the two randomly assigned mouse sperm groups. **f)** The binned (10 Mb) genetic distances plot along chromosome 1 for the two sperm groups. **g)** The bootstrapping distribution of the difference in total genetic distance between the two groups with two vertical lines indicating the lower bound and the upper bound of 95% confidence interval. **h)** Permutation results of difference in total genetic distance between two sperm cell groups with observed group difference indicated by the vertical line and p-value labelled on the top.

**Fig. 5.**
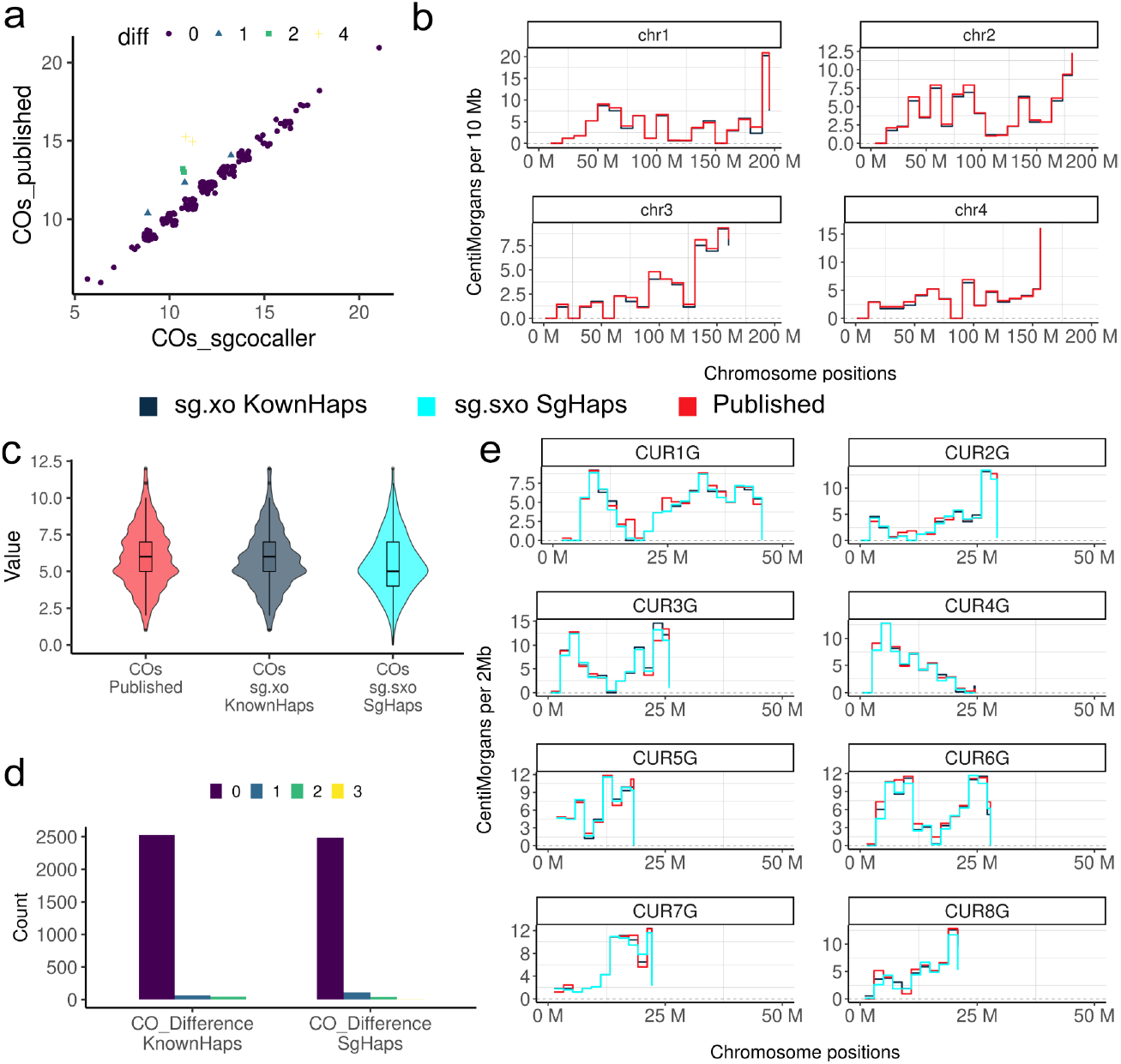
Comparisons between number of crossovers called from sgcocaller versus from the published study **a)** The number of crossovers (COs) called per sperm by sgcocaller (x-axis) versus by the original paper (y-axis) are plotted; points were jittered and colored and shaped by the differences in number of crossovers by the two methods. **b)** The genetic distances in 10 Mb chromosome bins were calculated and plotted along the chromosomes from using either crossover rates estimated from *sgcocaller xo* with the known haplotypes or the obtained published results of the mouse sperm dataset (21). **c)** The distribution of the number of crossovers called per apricot gamete (n = 329 apricot gametes) from the published study (20) (COs Published), calling *sgcocaller xo* using the published haplotype (sg.xo KnownHaps) and calling *sgcocaller sxo* with *sgcocaller phase* generated haplotype (sg.sxo SgHaps). **d)** The frequency of the difference in crossover counts with published crossover results per chromosome by sgcocaller using either the published haplotype (CO_Difference KnownHaps) or using *sgcocaller phase* generated haplotype (CO_Difference SgHaps). **e)** The genetic distances in every 2 Mb chromosome bin were calculated using crossover rates estimated from three approaches that showed high concordance along regions of the genome.

#### 10X-scCNV sequencing of gametes from a diploid apricot sample

We next applied sgcocaller and comapr on another example dataset of single-cell sequenced haploid pollen cells from an apricot tree generated using the 10X single-cell CNV protocol (20). The downloaded single-cell DNA reads from each apricot gamete (ENA:PRJEB37669) were re-aligned to the haplotype-resolved haploid genome (Currot) released from the same study. The hetSNP markers were identified by calling heterozygous variants from the pooled reads of all gametes *de novo* using `bcftools` and high-quality variants were kept (37, 44) (see ;Supplementary Methods).

Comparing with the crossover profile constructed from the original study, sgcocaller and comapr have generated a highly concordant crossover profile for the analysed gametes (Fig. 5c,d,e and Fig. S2). The number of crossovers identified per gamete (n = 329 gametes) was consistent with the published number of crossovers for these gametes by using both the published haplotype (sg.xo KnownHaps) and the *sgcocaller phase* generated haplotype (sg.sxo SgHaps) (Fig. 5c). Few chromosomes (4% by sg.xo KnownHaps; 5.6% by sg.sxo SgHaps) of the 2,632 chromosomes studied here exhibited differences in crossover counts (Fig. 5d). Out of the chromosomes that were called with different numbers of crossovers, 69% of them were called with fewer crossovers by sg.xo KnownHaps and 86% with fewer crossovers by sg.sxo SgHaps. Therefore, the discrepancy of results from the two methods resulted predominantly from sgcocaller being more conservative and calling fewer crossovers. Inspecting the crossover profile by chromosome regions has also demonstrated consistent genetic distances across the whole genome (Fig. 5e).

It is worth noting that not all cells available in the published study were analysed and included in the application results of sgcocaller due to the cell filtering step included in comapr which filtered out cells with excessive numbers of crossovers called and cells with poor SNP coverage. Chromosomes with excessive numbers of crossovers called are likely due to library preparation artefacts or abnormal chromosome segregation that results in implausibly many heterozygous SNPs in single gametes (Fig. S3). comapr offers a systematic way of filtering out these cases and can be applied to all gametes.

#### sgcocaller advances performance and efficiency

Many studies apply hidden Markov model based approaches for haplotype construction and crossover identification using sequencing or genotyping datasets (9, 21, 45–48). The majority of them, however, used customised or in-house scripts for crossover calling (9, 21, 45). Only a handful of them published reusable software tools or pipelines (40, 47), but none of them apply in the same scenarios as sgcocaller and comapr, which work directly on large-scale single-cell DNA sequencing datasets. The Hapi method, implemented in an R package, was proposed to construct chromosome-scale haplotypes using data from a small number of single gametes (40). Given a genotype matrix as input, it has functionalities overlapping with *sgcocaller phase* (40). We therefore compared the phasing performance of Hapi and *sgcocaller phase*|*swphase* on a total of four datasets: the individually sequenced mouse sperm cells (21), the 11 human sperm cells (the same dataset used in the performance evaluation of Hapi in the original paper (28, 40), the apricot gamete dataset generated by the 10X scCNV protocol (20), and lastly the subsampled mouse sperm dataset that is more comparable to a contemporary droplet-based single-gamete DNA sequencing dataset (Fig. 6e) (see Supplementary methods). To measure uncertainty in method performance, we constructed dataset replicates by dividing the original sets of gametes into ten portions and each time leaving out one portion (10%) of the gametes. Since there were only 11 cells in total in the human sperm cell dataset, 11 dataset replicates were constructed (n = 10 cells each). Ten distinct datasets were constructed from the apricot gametes (n = 330 or 331 gametes each), mouse sperm dataset (n = 174 or 175 sperm cells), and the subsampled mouse sperm dataset (n = 174 or 175 sperm cells).

**Fig. 6.**
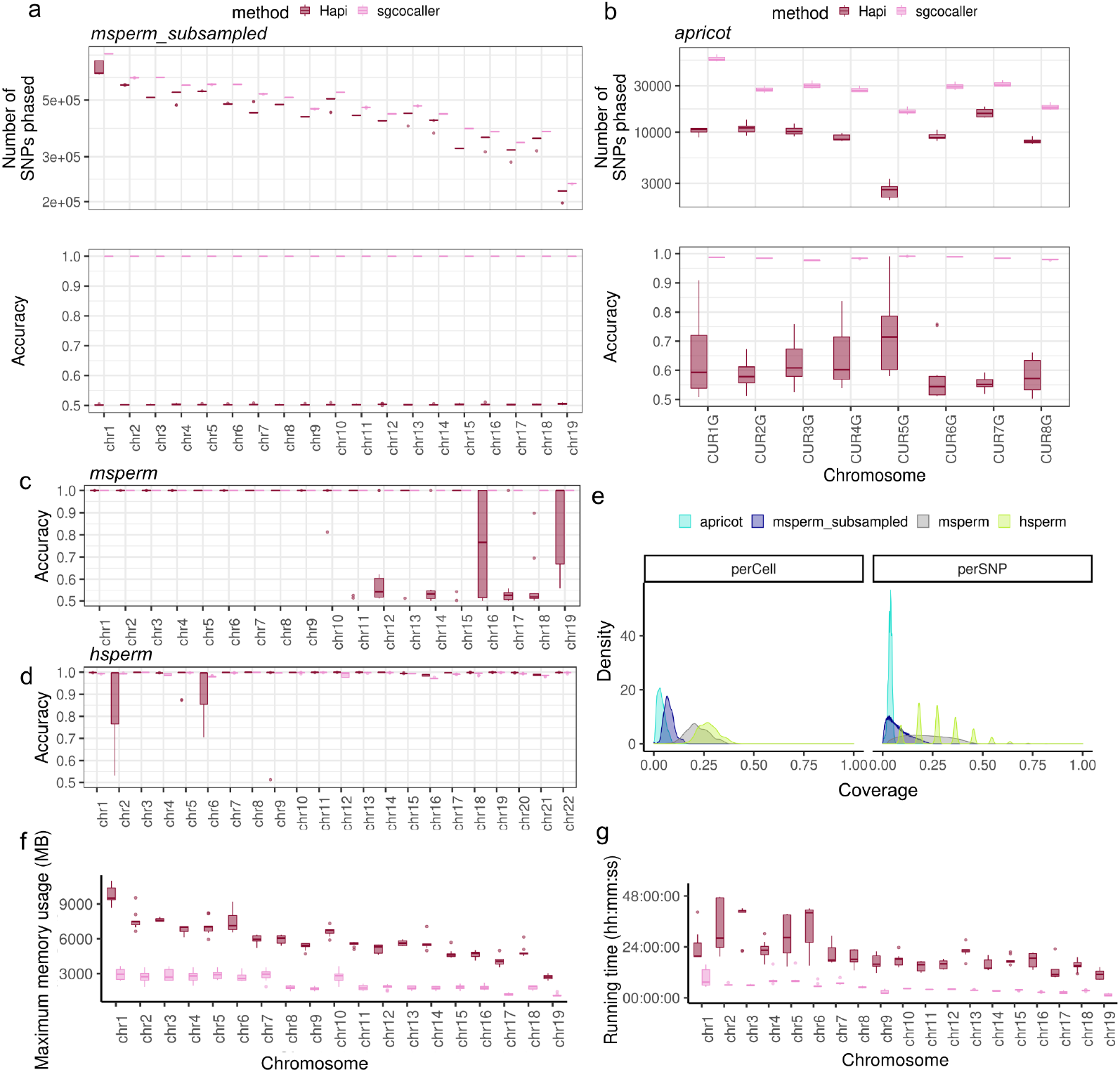
Phasing module performance comparison with existing method The phasing performance of sgcocaller and Hapi were compared on the further sub-sampled mouse sperm dataset (n = 194 sperm cells), apricot single-cell DNA sequencing dataset (n = 367 apricot gametes) (20), the mouse sperm sequencing dataset (n = 194 sperm cells) (21) and the human sperm sequencing dataset (n = 11 sperm cells) (28). **a)** The number of phased SNPs and phasing accuracy from running two methods (sgcocaller and Hapi) on 10 constructed datasets from using the further sub-sampled mouse sperm cells were compared (each with n = 174 or 175 sperm cells). **b)** Same as **a)** but for the constructed 10 apricot gamete datasets (each dataset with n = 330 or 331 gametes). Hapi failed when executing for two chromosomes and did not return values (2 out of 10 *×* 8 = 80 in total), which were not included in the plot. **c**,**d)** The phasing accuracy from running two methods on the 11 constructed datasets using 11 human sperm cells, and on the 10 constructed datasets using mouse sperm cells were plotted in boxplots. For human sperm dataset, Hapi failed to return values for 4 chromosomes (out of 11 *×* 22 = 242 in total), which were not included in the plots. **e)** The characteristics of the four datasets were compared by comparing the SNP coverage rate and cell coverage rate. **f**,**g)** The running time and memory usage by the two methods for phasing each chromosome using the constructed 10 datasets from the mouse sperm dataset (21) were compared. Time was reported in format of hour:minute:seconds, and memory was measured in units of mega bytes (MB). Measurements were reported by the ‘benchmark’ function from Snakemake (49).

We ran Hapi and *sgcocaller phase*|*swphase* on these constructed datasets. The phasing accuracy, measured by calculating the fraction of SNPs agreeing with the ground-truth haplotype sequence, and the number of phased SNPs by each approach were compared (Fig. 6a-d and Fig. S4).

On the lower coverage datasets, (the apricot gamete dataset and the sub-sampled mouse sperm dataset), the advantage of *sgcocaller phase*|*swphase* is readily apparent with far more SNPs phased and much higher accuracy. *sgcocaller phase*|*swphase* generated haplotypes that were concordant with the published haploid genome assembly with lowest accuracy across all chromosomes in all dataset repeats of 97.6% for the apricot dataset. All chromosomes were phased with accuracy above 99.99% for the sub-sampled mouse sperm datasets by *sgcocaller phase*|*swphase* (Fig. 6a.b). In contrast, Hapi delivered median accuracy below 75% for the apricot datasets and only 50% accuracy (equivalent to random guessing) for the sub-sampled mouse sperm datasets. Hapi’s poor accuracy on these datasets makes its results unusable for downstream applications like crossover calling.

On the high coverage datasets (the mouse sperm dataset and the human sperm dataset), *sgcocaller phase*|*swphase* achieved equivalent or greater haplotype completeness in terms of the number of phased SNPs compared to Hapi (Fig. S4). *sgcocaller phase*|*swphase* achieved near perfect accuracy for all sperm combinations (all above 0.99), whereas Hapi repeatedly generated unusable phasing results (accuracy *<* 0.6) for chromosome 12, 14, 17, 18 for the mouse sperm datasets (Fig. 6c). Hapi did not perform stably for the 11 human sperm datasets, but *sgcocaller phase*|*swphase* always generated phasing results with accuracy greater than 97% across all chromosomes in all human sperm combinations. (Fig. 6d). Hapi also failed for 4 chromosome runs that triggered errors (out of 242 chromosomes tested across all datasets), which were excluded from the comparison.

We compared the differences among the four datasets, which showed that the human sperm sequencing dataset and the mouse sperm sequencing data have much higher SNP coverage per cell and cell coverage per SNP compared to the apricot dataset (Fig. 6e and Fig. S5). The sub-sampled mouse sperm dataset has coverage distributions that are more comparable to the apricot dataset.

In summary, *sgcocaller phase*|*swphase* outperforms Hapi on both high coverage data and low coverage data with *sgcocaller phase*|*swphase* more dominant in the low coverage datasets.

#### Computational efficiency

We compared the computational efficiency of the two methods (sgcocaller and Hapi) on the large mouse sperm dataset and measured the running time as well as memory usage on each of the ten dataset repeats. The running time required by *sgcocaller phase* plus *sgcocaller swphase* was only 11% to 38% of the total time taken by Hapi (comparing the median running time of each chromosome). Considering *sgcocaller phase* timing results includes an extra step of parsing the DNA read file (BAM file) and variant file (VCF file), *sgcocaller phase* is even more advantageous in terms of speed than these results indicate. Examining diagnostic plots generated from *sgcocaller phase* can help with identifying switch errors so that *sgcocaller swphase* can be run selectively. Nevertheless, *swphase* was applied to all chromosomes and the total running times were still many-fold shorter than Hapi. In terms of memory usage, since *sgcocaller phase* and *sgcocaller swphase* were run consecutively, the process that used larger memory was regarded as the memory used by sgcocaller. From comparing the reported max_rss (the maximum amount of memory occupied by a process at any time that is held in main memory (RAM)) by `Snakemake` (49), sgcocaller required only 30% to 50% of the memory required by Hapi (Fig. 6f,g).

### Scalability of sgcocaller

We tested the scalability of different modules in sgcocaller with input files of varying sizes. Scalability testing was conducted by applying sgcocaller on chromosome 1 from the mouse sperm dataset. We measured the running time and memory usage of different modules by varying either the number of SNPs or the number of DNA reads available. When varying the number of DNA reads to process, the same input VCF which had 4.6 *×* 10^5^ SNPs was used, while the same BAM file with 1.1× 10^8^ reads was used when varying the number of SNPs. Running time and memory usage for all three modules scale linearly with the number of SNPs (nSNPs) to process. *sgcocaller sxo* runs significantly faster than *sgcocaller xo* as expected since it parses the allele counts matrices from *sgcocaller phase* instead of the input BAM and VCF file. Running time and memory usage of the three modules do not strictly increase with more DNA reads (nReads) to process, and plateau after nReads reaches about 320 million (Fig. S6). This performance is potentially due to the coverage rate of the available SNPs increasing as the number of DNA reads increases. However, when DNA reads are sufficient to cover all the available SNPs, increasing number of reads does not immediately increase running time or memory usage. Measurements were generated from running sgcocaller on a Linux server with `RedHat7.9(Maipo)` and with `–thread 4` for decompressing BAM input. The measurements of time and memory usage for different inputs were obtained using the “benchmark” function from the workflow manager `Snakemake` (49).

## Discussion

We have introduced a toolkit that consists of a commandline tool and a Bioconductor/R package for processing large-scale single-gamete DNA read datasets for individualised haplotype construction, crossover identification, and crossover landscape analysis. Personal haplotype construction is important in population genetics and clinical genetics for interpreting rare and disease-implicated variants (50, 51). Genomes of haploid gametes provide information of “longrange” haplotype blocks of the diploid donors and can be revealed via standard short-read sequencing. Single-gamete data generated using low-depth short-read sequencing are sufficient for reconstructing the two personal haplotypes by aggregating linkage information in a group of gametes.

Taking advantage of linkage information in single-gamete data, *sgcocaller phase* is able to construct personal haplotypes with standard short-read sequencing methods to near perfect accuracy. Comparisons with other phasing methods demonstrate that *sgcocaller phase* offers better performance on single-cell sequenced gametes in terms of accuracy and efficiency. Crucially, haplotypes constructed with *sgcocaller phase* solely from single-gamete sequencing data are sufficiently accurate to support highly accurate downstream crossover calling with sgcocaller’s crossover calling methods. sgcocaller achieves highly accurate phasing results in settings where competitors fail to generate usable haplotypes for crossover calling. The outputs of sgcocaller can be conveniently imported into R with the comapr package for sophisticated crossover landscape visualisation and analysis. The application demonstrations of *sgcocaller*|*comapr* on public datasets show that *sgcocaller*|*comapr* are able to produce stable and higher accuracy results than other methods with greater convenience and computational efficiency for large datasets. Although *sgcocaller*|*comapr* are optimised to work with cell-barcoded DNA reads, we also demonstrated and showed examples of how our tools can be applied on bulk-sequenced individual gametes. Using the modern programming language Nim and building on top of C-based libraries, sgcocaller processes the large genomic datasets efficiently. The multiple release formats of sgcocaller enable it to be more accessible to the community.

Advances in single-cell DNA sequencing technologies that generate larger-scale and higher-quality datasets enable exciting opportunities for research using personalised meiotic crossover landscapes (11). Abnormal meiotic crossovers are often associated with infertility (5). Therefore, characterising individual meiotic crossover profiles and studying crossovers as a phenotype using single-cell techniques has application in reproduction clinics. Fundamental mechanisms and factors affecting meiotic crossovers remain active research topics. Using single-gamete methods for generating and comparing crossover profiles of individuals with different conditions has many advantages over traditional approaches that require recruiting large sample sizes (11). Our software serves as a complete toolkit for the first step of studies to inspect the single gamete data of an individual for understanding individual-level variations in meiotic crossovers.

Due to the current design of the methods, *sgcocaller*|*comapr* are limited to the analysis of gametes from diploid organisms. Currently, sgcocaller only supports using multiple threads for decompressing the input BAM file and recommends users to run each chromosome in parallel to reduce computational time. Future work in the development of sgcocaller might include updating the program to include the multi-threading feature by chromosomes internally and supporting analyses for polyploid organisms.

## Conclusion

Available in multiple release formats (nimble package, static binary, docker image, bioconda package), sgcocaller fills the gap of a highly accurate and efficient tool with simple installation for constructing personalised haplotypes and calling crossovers in single gametes from single-cell and bulk DNA sequenced gametes. The availability of the companion Bioconductor/R package comapr enables the downstream statistical analysis of crossovers in the cells and among individuals, integrating well with existing R packages for various visualisation and exploratory analysis tasks. In concert, these packages represent a comprehensive, user-friendly toolkit for the construction and analysis of personalised crossover landscapes from single-gamete sequencing data.

## Availability of data and materials

Source code of the latest version of sgcocaller is publicly available at a GitLab repository and comapr at a GitHub repository under a MIT license. comapr is also available as a Bioconductor package. The datasets used in this study are public datasets with raw sequencing data downloaded from GEO with accession GSE125326 and ENA with project ID PRJEB37669. The analysis steps including preprocessing and scripts for generating the figures are openly accessible at GitLab repositories and workflowr (52) web pages demonstrating the full workflow of applying *sgcocaller*|*comapr* are available. The source code and analysis repositories included in this study are listed below:

1. sgcocaller: https://gitlab.svi.edu.au/biocellgen-public/sgcocaller
2. comapr: https://github.com/ruqianl/comapr and https://bioconductor.org/packages/comapr
3. Mouse sperm data analysis: https://gitlab.svi.edu.au/biocellgen-public/hinch-single-sperm-DNA-seq-processing
4. Apricot gamete analysis: https://gitlab.svi.edu.au/biocellgen-public/calling-crossover-from-single-gamete-sequencing-of-apricot
5. appendCB: https://github.com/ruqianl/appendCB
6. distributions: https://github.com/ruqianl/distributions

## Acknowledgements

The authors thank Brendan Hill for reviewing the source code of sgcocaller. The double Holliday junction in the comapr Hex Logo was made with affinity designer and Biorender.

## Supplementary Note 1: Supplementary materials

Available online

