## Supplementary materials for "sgcocaller and comapr: personalised haplotype assembly and comparative crossover map analysis using single-gamete sequencing data"

April 22, 2022

[1]Bioinformatics and Cellular Genomics, St. Vincent's Institute of Medical Research,9 Princes Street,Fitzroy Victoria, 3065, Australia [2]Melbourne Integrative Genomics/School of Mathematics and Statistics, Faculty of Science, The University of Melbourne, Building 184, Royal Parade, Parkville, Victoria,3010, Australia [3]DNA Repair and Recombination Laboratory, St Vincent's Institute of Medical Research,9 Princes Street,Fitzroy Victoria, 3065 Australia [4]The Faculty of Medicine, Dentistry and Health Science, The University of Melbourne, Victoria 3010 Australia

**5 Calling crossovers** **11**

**6 Supplementary figures** **13**

### 1 sgcocaller phase

Haploid genomes of the pool of gametes collected from an individual are the results of meiosis and meiotic crossovers. Based on the known mechanisms of meiosis and crossovers, with respect to each chromosome half of the gametes produced in meiosis will not have crossovers and thus they represent the original (maternal/paternal) haplotypes of the diploid donor. Thus, for a given chromosome, we expect the rest of gametes to have crossovers (i.e., the chromosomes are formed by the combined parental haplotypes of the donor). Crossovers are low frequency events across chromosomes and crossover positions are sparse. The SNP linkages in small chromosome regions across all haploid gametes are therefore reliable for reconstructing the donor's haplotypes.

#### 1.1 Single gamete genotype matrix

To generate the two haplotypes of each chromosome for the diploid donor from gametes, the first step is finding the list of unphased hetSNPs (heterozygous SNP loci that differ between the maternal and paternal homologous chromosomes) by standard variant calling tools such as `bcftools` using pooled DNA reads from gametes. Only biallelic SNPs are considered in this step, therefore finding one haplotype also implicitly resolves the second haplotype by switching all alleles to its alternative at each hetSNP. In other words, the two haplotypes are bitwise complementary to each other. With the hetSNPs known, the DNA reads from each gamete are parsed and summarised into a genotype matrix for each chromosome with values of 1 or 2 corresponding to matching with the REF allele and the ALT allele at each unphased hetSNP. ALT allele read frequency (AF) is used for genotyping each SNP in each gamete and AF $\leq 0.3$ is genotyped as 1 (REF) while SNPs with AF $\geq 0.7$ are genotyped as 2 (ALT). Filtering options are available for excluding low mapping/base quality reads, low coverage SNPs per cell with the options *–minCellDP* and *–minTotalDP*.

#### 1.2 Template cell identification

*sgcocaller phase* first attempts to identify a gamete cell without crossovers as a template cell whose sequence of genotypes are used as the initial inferred haplotype. An ideal template cell for phasing each chromosome is a cell without crossovers and having sufficient SNP coverage regarding this chromosome. To find a cell without crossovers, *sgcocaller phase* finds three pairs of gametes with lowest genotype dissimilarity. The dissimilarity of two genotype sequences is calculated by finding the proportion ($p$) of disconcordant SNPs between the two sequences and we define the dissimilarity value as $min(p, 1-p)$. *sgcocaller phase* includes $maxDissim$ as a user supplied option that controls how similar ($1-maxDissim$) it requires for two cells to be considered as template cells. *sgcocaller phase* finds a maximum of three cell pairs as potential template cells. When multiple template cell pairs are available, the cell with the highest SNP coverage is chosen as the template cell. This approach is based on the idea that when two gamete chromosomes have no crossovers, either their genotype sequences will be the same (or at least very similar if the same parental haplotypes have been inherited) or totally different (when different parental haplotypes are inherited by the two chromosomes). When multiple template cell pairs are available, the cell with the highest SNP coverage is chosen as the template cell.

There are rare cases when two gametes have crossovers at exactly the same positions, which also leads to high genotype similarity in the gametes. Such cases can be revealed by diagnostic plots (Fig. 2c,d) with the simple R script provided and then corrected by *sgcocaller swphase*. The template cell to use can also be chosen manually by the user via the *templateCell* option.

#### 1.3 Infer missing SNPs in the haplotype template

Upon forming a haplotype template, *sgcocaller phase* increases the completeness of the haplotype template by inferring the genotype of the missing SNPs (that is, SNPs with no read coverage) from the template using other gametes in which the SNP is available (that is, has read coverage). A haplotype template represents two actual haplotype (allele) sequences $(h, h')$ that are bitwise complementary with respect to the REF and ALT alleles defined for the hetSNPs used; one haplotype is thus derived by changing all alleles from the complementary haplotype to their alternatives. Intuitively, we want to use linkage information from other gametes to "fill in the gaps" in the template haplotype by finding gametes with coverage at the missing SNP site. Loosely, if a gamete with coverage at the missing SNP has the same haplotype as the template, then we assume that the template should have the same allele at the missing SNP as the gamete with coverage. If the gamete with coverage has the complementary haplotype, then the missing SNP in the template should have the alternative allele.

More precisely, to infer missing SNPs' genotypes in the haplotype template, we need to find the linkage type of the SNP to the haplotype template. Only two types of linkages are possible due to the existence of two possible alleles (e.g., 1 or 2 ) for a given SNP. The first linkage type ("type 1") refers to gametes with the template haplotype and allele 1 at the missing SNP and gametes with the haplotype complementary to the template haplotype and allele 2 at the missing SNP; that is, the missing SNP should have allele 1 in the template (type 1:$\{(1, h), (2, h')\}$). The second linkage type ("type 2") is the inverse, such that the template should have allele 2 at the missing SNP (type 2:$\{(1, h'), (2, h)\}$). To determine the linkage type of the SNP in supporting gametes (that is, those with read coverage of the SNP), the nearby SNPs' genotype sequence in each supporting gamete (specifically, the 10 closest SNPs with read coverage in both the template gamete and supporting gamete) is first compared with the template haplotype at the matching positions to define the haplotype (template or complementary) of the supporting gamete. With the haplotype of the supporting gamete determined, the linkage type supported by the gamete immediately follows. The posterior probabilities of linkage types of a missing SNP are calculated by looking at the number of gametes supporting each type of SNP linkage. Assuming a genotype error rate of 0.1 and that the two linkage types are equally likely to happen, the posterior probability of each linkage type of a missing SNP can be calculated:

$t_1$: {type 1 linkage counts}    $t_2$: {type 2 linkage counts}

$$\begin{aligned}
p(\text{type } 1|t_1, t_2) & \\
&= \frac{p(t_1,t_2|\text{type } 1)p(\text{type } 1)}{p(t_1,t_2|\text{type } 1)p(\text{type } 1)+p(t_1,t_2|\text{type } 2)p(\text{type } 2)} \\
&= \frac{p(t_1,t_2|\text{type } 1)}{p(t_1,t_2|\text{type } 1)+p(t_1,t_2|\text{type } 2)} \\
&= \frac{0.9^{t_1}0.1^{t_2}}{0.9^{t_1}0.1^{t_2}+0.1^{t_1}0.9^{t_2}} \\
p(\text{type } 2|t_1, t_2) &= \frac{0.9^{t_2}0.1^{t_1}}{0.9^{t_1}0.1^{t_2}+0.1^{t_1}0.9^{t_2}}.
\end{aligned}$$

We use a threshold cut-off (default is 0.99 and it can be changed via the option *posteriorProbMin*) for determining whether a missing SNP can be inferred. Missing SNPs with posterior probabilities over the threshold are inferred to be the suggested linkage type. Applying this approach genome-wide, we can make maximal use of read coverage across all gametes to maximise the completeness of the template haplotype. After inferring missing SNPs from the template haplotype, the step of inferring SNPs is then performed against all hetSNPs in the template haplotype to correct any genotyping errors in the template haplotype.

### 2 sgcocaller swphase

When an ideal template cell is not used for phasing in the previous step, the chosen template cell may have crossovers leading to switching errors in the inferred haplotype (Fig 2c). *sgcocaller swphase* is able to detect the switch errors and generate the corrected haplotype. To save unnecessary computing, *sgcocaller swphase* calculates the switch scores only for identified SNP bins whose positions have high risk of having switch errors. High risk SNP bins are found by firstly grouping all hetSNPs into bins of 2,000 consecutive SNPs with a moving step of 200 SNPs (both are changeable via options when running *sgcocaller swphase*). The proportion of gametes having crossovers are calculated for each bin. A SNP bin is labelled as a high risk bin when the proportion of gametes having crossovers is above 0.5. *sgcocaller swphase* calculates switch scores for SNPs positions potentially having switch errors. It is based on the idea that when the majority of gametes have crossovers according to the template haplotype, it indicates a "crossover" or switch error in the template haplotype. The crossover identification in this step is fast as it simply compares the dissimilarity of gametes' genotype sequences with the inferred haplotype sequence for the SNPs in each bin. A default threshold value of 0.0099 is set for the dissimilarity to decide whether a crossover has happened in the gamete or not.

#### 2.1 Switch score calculation

To construct the switch score (formed using a concept for splitting blocks similar to that from a previous haplotype construction method [1]), which represents how likely a switch error has happened at SNP $i$ in the inferred haplotype, the switched haplotype is first constructed. Let $H_l^i$ represents the haplotype sequence to the left $N$ bases of SNP $i$, and $H_r^i$ represents the haplotype sequence to right $N$ bases of SNP $i$. The current haplotype around SNP $i$ is $H^i = \{H_l^i, H_r^i\}$. The switched haplotype is a new sequence of $H_{sw}^i = \{H_l^i, H_r^{i\prime}\}$, where $H_r^{i\prime}$ is the bitwise complementary sequence of $H_r^i$. The switch score for each SNP $i$ is calculated using the log-ratio of the probability of observing the genotype sequences in all gametes given the switched haplotype with the probability of observing the genotype sequences in all gametes given the non-switched haplotype. Assuming the occurrence of a switch error or not at any site is random, that is the prior of having switch or no switch at a SNP site equals 0.5, the switch score is the log-ratio of the posterior probabilities of the switched haplotype and the not-switched haplotype. When calculating the probability of observing each gamete's genotype sequence given the haplotype ($H^i$ or $H^i_{sw}$), only the local ($N$) SNPs are considered (controlled by the *lookBeyondSNPs* option with default set to 20).

The haplotype $H^i$ is a sequence of alleles and also implicitly defines the second haplotype $H^{i\prime}$ which can be derived by switching all alleles to their complementary alleles (in the called genotypes of the hetSNPs). The probability of observing all gametes' genotype sequences at the local ($N$) SNPs around SNP $i$ given the local haplotype $(H^i, H^{i\prime})$ is,

$$p(G^i \mid H^i, H^{i\prime}) = \prod_{j=1}^{m} p(G^i_j \mid H^i, H^{i\prime}),$$

where $G^i$ represents local genotypes around SNP $i$ of all ($m$) gametes, and $G^i_j$ represents local genotypes at SNP $i$ from gamete $j$. For each gamete $j$, the probability of its genotype sequence given $(H^i, H^{i\prime})$ is

$$p(G^i_j \mid H^i, H^{i\prime}) = \frac{p(G^i_j \mid H^i) + p(G^i_j \mid H^{i\prime})}{2}.$$

Similarly for the switched haplotype $H^i_{sw}$, the probability of observing the local genotypes of all gametes given

$$(H^i_{sw}, H^{i\prime}_{sw}),$$

$$p(G^i \mid H^i_{sw}, H^{i\prime}_{sw}) = \prod_{j=1}^{m} p(G^i_j \mid H^i_{sw}, H^{i\prime}_{sw}),$$

where

$$p(G^i_j \mid H^i_{sw}, H^{i\prime}_{sw}) = \frac{p(G^i_j \mid H^i_{sw}) + p(G^i_j \mid H^{i\prime}_{sw})}{2}.$$

In addition, probability of observing a gamete's genotype sequence given a haplotype sequence, $p(G_j \mid h)$, is calculated as (assuming genotype error rate 0.1 and using $d$ as the number of different bases between the allele sequence in $G_j$ and the allele sequence in haplotype allele sequence $h$):

$$p(G_j \mid h) = 0.1^d \times 0.9^{(K-d)},$$

where $K$ is the number of co-existing SNPs in two allele sequences under comparisons. The switch score for SNP $i$ is thus

$$S_i = \log \frac{p(G^i \mid H^i_{sw}, H^{i\prime}_{sw})}{p(G^i \mid H^i, H^{i\prime})}.$$

Upon calculating switch scores for a sequence of SNP positions, the switching point is identified as the peak of a stretch of positive switch scores (Fig. 2b).

The minimum threshold for identifying switch points is set via the option *minSwitchScore*, which can vary depending on features of the dataset including the number of cells available. The template haplotype is then corrected by flipping all SNP alleles to their complementary alleles after an identified switch point, thus generating the corrected haplotype with the switch error removed.

### sgcocaller xo and sgcocaller sxo

Crossovers can be detected by finding haplotype shifts in the gametes' haploid genomes. With DNA reads from each cell mapped, the haplotypes of SNP markers are inferred by looking at the alleles carried by the DNA reads mapped to genomic positions of these SNP markers. However, with technical artefacts (from sequencing and mapping), it is expected to observe some proportion of conflicting alleles from the underlying haplotype (Fig. 1b). To reconstruct the haplotype structure of the haploid genomes from the mapped DNA reads while accounting for technical noise including mapping errors for crossover identification, *sgcocaller xo* applies a Hidden Markov model (HMM) with a binomial emission model (Fig. 2e).

The two hidden states in the HMM represent the haplotype origins of DNA segments (represented by allele types of a list of SNPs) in the gametes' genomes. Transitioning from one state to another between two SNP markers corresponds to a crossover detected and the transition probability is set to be dependent on the two SNP markers' base pair distances (physical distances) [2]. The transition probability is programmed as an configurable option (*cmPmb*) in *sgcocaller xo*. The relationship between observed allele counts and the underlying hidden states are modelled by the two binomial distributions whose success rates are also user-configurable options (*–thetaREF, –thetaALT*) in *sgcocaller xo* (Fig. 2e and MATERIALS AND METHODS ). The two states (named as REF and ALT) match with the alleles (bases) in the REF and ALT fields in the input VCF. The binomial distributions in the HMM model the ALT allele read counts at each SNP site and thus the *–thetaREF* value is expected to be small (e.g., 0.1) while *–thetaALT* value is expected to be higher (e.g., 0.9).

#### 2.2 The Hidden Markov Model

The two hidden states in the HMM represent the haplotype origins of DNA segments (represented by allele types of a list of SNPs) in the gametes' genomes. Transitioning from one state to another between two SNP markers corresponds to a crossover detected and the transition probability is set to be dependent on the two SNP markers' base pair distances (physical distances) [2]. The transition probability is programmed as an configurable option (*cmPmb*) in *sgcocaller xo*. The relationship between observed allele counts and the underlying hidden states are modelled by the two binomial distributions whose success rates are also user-configurable options (*–thetaREF, –thetaALT*) in *sgcocaller xo* (Fig. 2e and MATERIALS AND METHODS). The two states (named as REF and ALT) match with the alleles (bases) in the REF and ALT fields in the input VCF. The binomial distributions in the HMM model the ALT allele read counts at each SNP site and thus the *–thetaREF* value is expected to be small (e.g., 0.1) while *–thetaALT* value is expected to be higher (e.g., 0.9). The data in this model are the allele-specific read counts across the list of hetSNP sites for each chromosome, whereas the underlying haplotype of a SNP site on the chromosome is a hidden variable to be inferred. In the case of gametes, which have haploid genomes, there are two possible hidden states for each SNP corresponding to the two haplotypes of the parent. The two possible alleles at each hetSNP site can be referred as REF or ALT arbitrarily. The REF or ALT state for each SNP also aligns with the same REF or ALT allele in the provided VCF input file. At each SNP site $i$, the two hidden states: $s^i = \text{alt}$ corresponds to ALT haplotype while $s^i = \text{ref}$ corresponds to REF haplotype. The emission probabilities are modelled by two binomial distributions:

$$c^i_{\text{total}} = c^i_{\text{alt}} + c^i_{\text{ref}} \ ,$$

$$c^i_{\text{alt}} | s^i = \text{alt} \sim \text{Bin}(c^i_{\text{total}}, p_{\text{alt}}) \ ,$$

$$c^i_{\text{alt}} | s^i = \text{ref} \sim \text{Bin}(c^i_{\text{total}}, p_{\text{ref}}) \ ,$$

where $c^i_{\text{alt}}$ and $c^i_{\text{ref}}$ denote the alternative allele read count and reference allele read count at SNP $i$, respectively, $c^i_{\text{total}}$ denotes the total read count at SNP $i$, and $p_{\text{alt}}$ and $p_{\text{ref}}$ are configurable parameters when running *sgcocaller xo* and denotes the success rates in the two binomial distributions respectively. The transition probabilities ($p^i_{\text{trans}}$) are modelled dependent on markers' base pair distances [2] with the default of average 0.1 centiMorgan per 1Mb (1 million base pairs) which can be changed via the *cmPmb* option. Lastly, the initial probabilities for the two hidden states are both set to be 0.5, making the assumption that they are equally likely to happen.

#### Measuring support for detected crossovers

We use a quantitative measure, the log-likelihood ratio (logllRatio), for measuring the amount of support from data for detected crossovers. We define a (Viterbi) state segment as a consecutive list of SNPs with the same state. We also use the term "inferred state" to refer to the state inferred through applying the Viterbi algorithm, whereas we use the term "altered state" of a SNP to refer to the state obtained by altering its inferred state to its opposite. We measure the support in the data for the detected crossover using the log-likelihood ratio for the segment with the inferred state, relative to the altered state.

Specifically, the logllRatio is calculated by taking the log-likelihood of the data given the current inferred state minus the log-likelihood of the data given the altered state (Fig. 2f)

##### 2.2.1 logllRatio calculation

For a state segment that spans SNP $q$ to $k$, assuming the state segment has been inferred with state $REF$,

$$\text{Logll}_{\text{inferred}} = \log(t_l) + \sum_{i=q}^{k} \log f(c^i_{\text{alt}}; c^i_{\text{total}}, p_{\text{ref}}) + \log(t_r),$$

where $t_l$ and $t_r$ denote the transition probabilities from SNP $q-1$ to $q$ and from SNP $k$ to $k+1$, respectively, and $f$ is the binomial probability mass function given by

$$f(x; c, p) = \binom{c}{x} x^p (c - x)^{1-p}.$$

The log-likelihood of the data given the altered state is thus

$$\text{Logll}_{\text{altered}} = \log(1 - t_l) + \sum_{i=m}^{k} \log f(c^i_{\text{alt}}; c^i_{\text{total}}, p_{\text{alt}}) + \log(1 - t_r).$$

The logllRatio is derived by

$$\text{Logll}_{\text{inferred}} - \text{Logll}_{\text{altered}}.$$

##### 2.3 The outputs

Due to the sparsity of single-cell datasets, we have used sparse matrices in Matrix Market format as the output for *sgcocaller xo*, which saves disk space and enables efficient parsing in downstream analysis. *sgcocaller xo* generates sparse matrices of the allele counts, inferred haplotype states (output of the Viterbi algorithm) for each SNP across each cell. Columns of these matrices correspond to the list of single gamete cell barcodes and rows correspond to the list of het-SNPs. A supplementary text file (*viSegInfo.txt*), which contains the summary features of inferred Viterbi segments (a list of consecutive SNP markers with the same inferred haplotype state) (Fig. 2f), is also provided and can be used for convenient post-processing such as filtering of false positive crossovers. The haplotypes of each SNP are inferred using the Viterbi algorithm [3] therefore the states/segments are also referred to as the Viterbi state/segments. Features including starting SNP position, ending SNP position, the number of SNPs supporting the segment, and the log-likelihood ratio of the Viterbi segment are recorded for each segment in the text file (*viSegInfo.txt*).

#### 3 comapr - resampling-based methods

To test for differences in the number of crossovers between any two groups of cells, re-sampling methods, permutation and bootstrapping tests [4, 5, 6], have been implemented in *comapr*. The two resampling-based functions in *comapr* are able to either generate the to generate the empirical p-value or find the confidence intervals for the estimate of the group differences.

The `permuteDist` function performs permutation:

1. Record the observed difference in total genetic distances between the two groups of cells, $d_{obs}$.

2. Take the group labels vector and permute the group labels by randomly assigning the labels across cells and calculate the new difference with the newly-generated permuted grouping.

3. Repeat step 2 for $B$ times (e.g., $B = 1,000$).

4. Calculate the permutation p-value by using the `permp` function from the `statmod` package [7] that calculates the appropriate p-values for the permutation test when permutations are sampled with replacement, avoiding the common pitfalls of under-estimated p-values [4, 7].

The steps for generating the bootstrapping confidence intervals of genetic distance differences in two groups (A and B) of cells are implemented in the `bootstrapDist` function:

1. Randomly draw $n$ cells with replacement from cells in group A where $n$ is the group size of A and calculate the total genetic distance $d_1$ with the cells drawn.

2. Randomly draw $m$ cells with replacement from cells in group B where $m$ is the group size of B and calculate the total genetic distance with the cells drawn $d_2$.

3. Calculate difference in total genetic distances by $d_1 - d_2$.

4. Repeat step 1-3 for $B$ times (e.g., $B = 1,000$).

5. Calculate bootstrap confidence intervals for the sampling distribution of the difference in genetic distance using the `quantile` functions at a desired level from R [8].

Whether or not the acquired confidence intervals contain zero can be use to decide whether or not the difference is statistically significant at a desired level.

### 4 Phasing performance comparison

The detailed function calls for applying the two methods on the constructed datasets using human sperm, apricot gametes and mouse sperm datasets are described below.

#### 4.1 Human sperm cell dataset

Hapi was run following the tutorial example and the allowNA parameter in imputing missing genotypes function was set to 3. *sgcocaller phase* was run using options: `--threads 4 --barcodeTag CB --minDP 2 --maxDP 50 --maxTotalDP 200 --minTotalDP 6 --maxDissim 0.099--minSNPdepth 1 --maxExpand 1000 --posteriorProbMin 0.9` for each chromosome in each dataset. *sgcocaller swphase* was called for all chromosomes with `--lookBeyondSnps 150 --minSwitchScore 580` and `--minPositiveSwitchScores 100`.

#### 4.2 10X scCNV apricot gametes

Due to the sparsity of the dataset, the `allowNA` argument in the `hapiImupte` function of Hapi was raised to 310 to avoid triggering running errors. *sgcocaller phase* was applied with options `--threads 4 --barcodeTag CB --minDP 2 --maxDP 10 --maxTotalDP 80 --minTotalDP 6 --minSNPdepth 1 --posteriorProbMin 0.99` and *sgcocaller swphase* was called with `--binSize 1000 --stepSize 800`.

##### 4.3 Mouse sperm dataset

For the mouse sperm dataset [9], 10 datasets were constructed by leaving out a different 10% of the gametes each time. The phase of the known mouse strains were used as the ground truth. The alleles in the list of called hetSNPs was swapped for every other position to create the list of unphased hetSNPs. The `allowNA` argument in the `hapiImupte` function of Hapi was set to 3 to avoid triggering running errors. *sgcocaller phase* was applied with options `--threads 4 --barcodeTag CB --minDP 2 --maxTotalDP 350 --maxDP 10 --minSNPde pth 5 --maxDissim 0.0099 --posteriorProbMin 0.99` and *sgcocaller swphase* was called with `--binSize 1000 --stepSize 800 --lookBeyondSnps 10` default settings.

For the further sub-sampled mouse sperm dataset, the allowNA was set to be 3 and *sgcocaller phase* was called with options `--threads 4 --barcodeTag CB --chrom wildcards.chr --minDP 2 --maxTotalDP 150 --maxDP 10 --minSNPde pth 1 --maxDissim 0.0099`. *sgcocaller swphase* was called with options `--binSize 1000 --stepSize 800 --lookBeyondSnps 10`.

#### 5 Calling crossovers

##### 5.1 Mouse sperm dataset

Crossover results for mouse sperm data were obtained from calling *sgcocaller xo* on the prepared phased hetSNPs and BAM file for 194 sperm cells as described before (see MATERIALS AND METHODS). The detailed code available in a public GitLab repository (see AVAILABILITY OF DATA AND MATERIALS). with following filtering settings. *sgcocaller xo* was called with options: `--maxTotalDP 450 --maxDP 10 --thetaREF 0.1 --thetaALT 0.9`. The final crossover intervals were identified using *comapr* with detailed code available in a public GitLab repository (see AVAILABILITY OF DATA AND MATERIALS) with following filtering settings.

Segment level filters:

1. minSNP=30, the segment that results in one/two crossovers has to have more than 30 SNPs of support.

2. minlogllRatio=150, the segment that results in one/two crossovers has to have logllRatio larger than 150.

3. bpDist=$10^5$, the segment that results in one/two crossovers has to have base pair distances larger than $10^5$.

Cell level filters:

1. maxRawCO=55, the maximum number of raw crossovers allowed on a chromosome (excessive number of crossovers indicate abnormal chromosomes. Cells with any chromosome having more crossovers than the *maxRawCO* are be excluded.

2. minCellSNP=200, there has to be more than 200 SNPs detected within a cell per chromosome, otherwise this cell is removed.

#### 5.2 10X scCNV apricot data with known haplotype

*sgcocaller xo* was applied on 367 apricot gametes to identify crossovers using the hetSNPs called (see Section Preprocessing public datasets) with option: `--thetaREF 0.3 --thetaALT 0.7 --maxTotalDP 280 --maxDP 30 --minTotalDP 20 --minDP 2 --cmPmb 1e-12`. The following fitering thresholds were applied when parsing and interpreting crossover intervals from *xo*'s results: minSNP=10, minlogllRatio=5, maxRawCO = 5, minCellSNP=200, bpDist=$10^6$. `bpDist` was set to $10^5$ for chromosome 1 and scaled by relative chromosome size for the rest of the chromosomes.

#### 5.3 10X scCNV apricot data with sgcocaller phased haplotype

*sgcocaller sxo* was applied on *sgcocaller* phased haplotypes and used the generated output files from the previous step, thus avoiding redundant processing including parsing DNA alignment files. The following options were applied `--minTotalDP 5 --minDP 1 --thetaREF 0.1 --thetaALT 0.9 --cmPmb 1e-9 --maxDP 6 --maxTotalDP 50`. The following fitering thresholds were applied when parsing and interpreting crossover intervals from *sxo*'s results: minSNP = 5, minCellSNP = 80, maxRawCO = 5, minlogllRatio = 0, bpDist=$10^6$. `bpDist` was set to $10^5$ for chromosome 1 and scaled by relative chromosome size for the rest of the chromosomes.

### 6 Supplementary figures

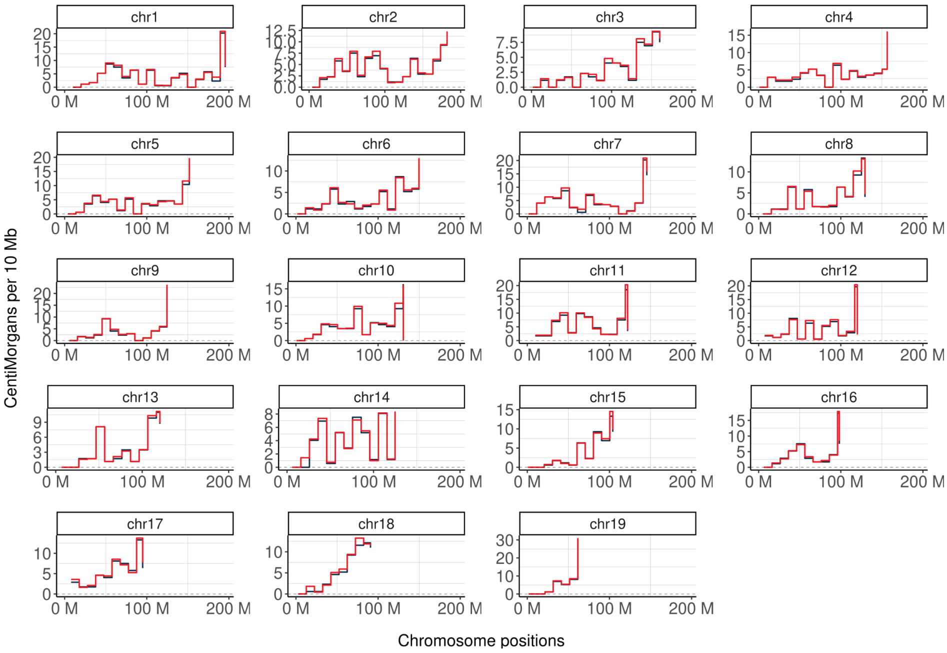

Figure S1: Genetic distances calculated by the *sgcocaller xo* and the published results from the original study in 10 Mb chromosome bins on the mouse sperm dataset for all autosomes. The same plot as in Fig5 b.

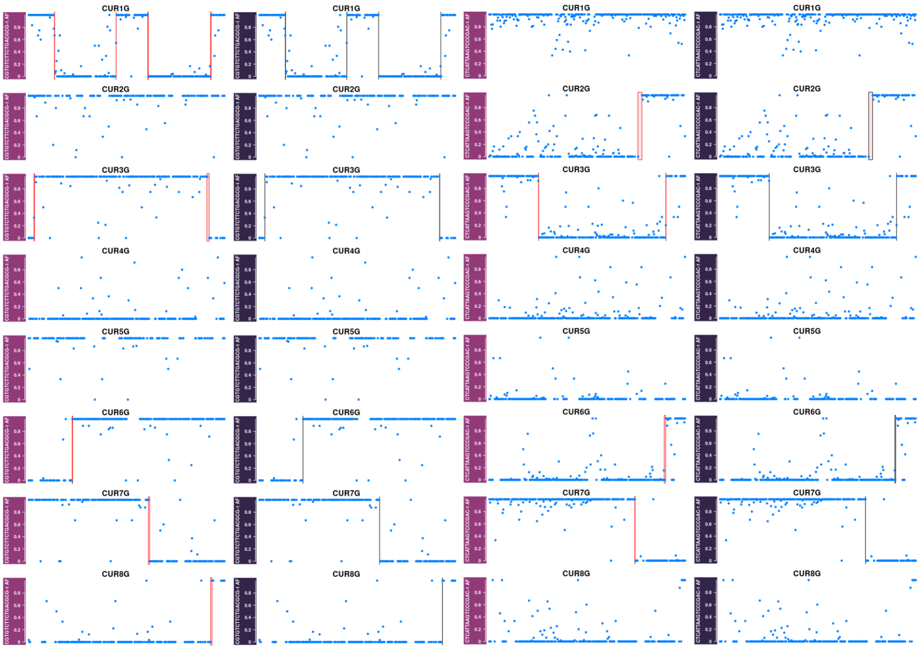

Figure S2: Crossovers called by sgcocaller-comapr workflow(left columns in two panels) and from published study (right columns in two panels) for two randomly selected apricot pollen

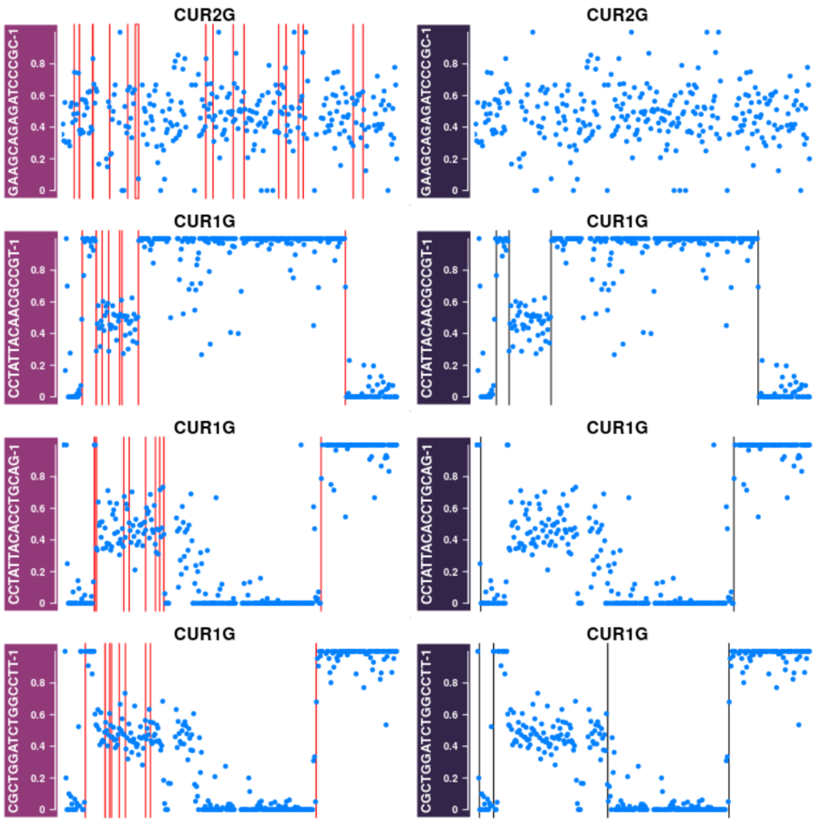

Figure S3: Examples of chromosomes with excessive crossovers called by *sgco-caller xo* and filtered in analysis (left column). Right column plots the crossover results released by previous study [9]

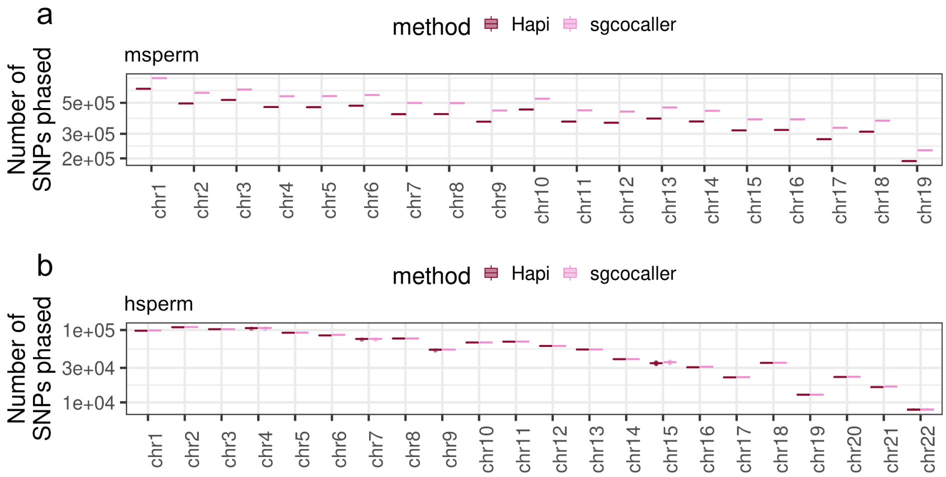

Figure S4: Number of phased SNPs by the two methods on two different datasets **a)** The number of phased SNPs by the two methods on the constructed ten mouse sperm datasets from using the original set of mouse sperms [2]. **b)** The number of phased SNPs by the two methods on the constructed eleven human sperm sperm datasets from using the original 11 human sperm cells [10]

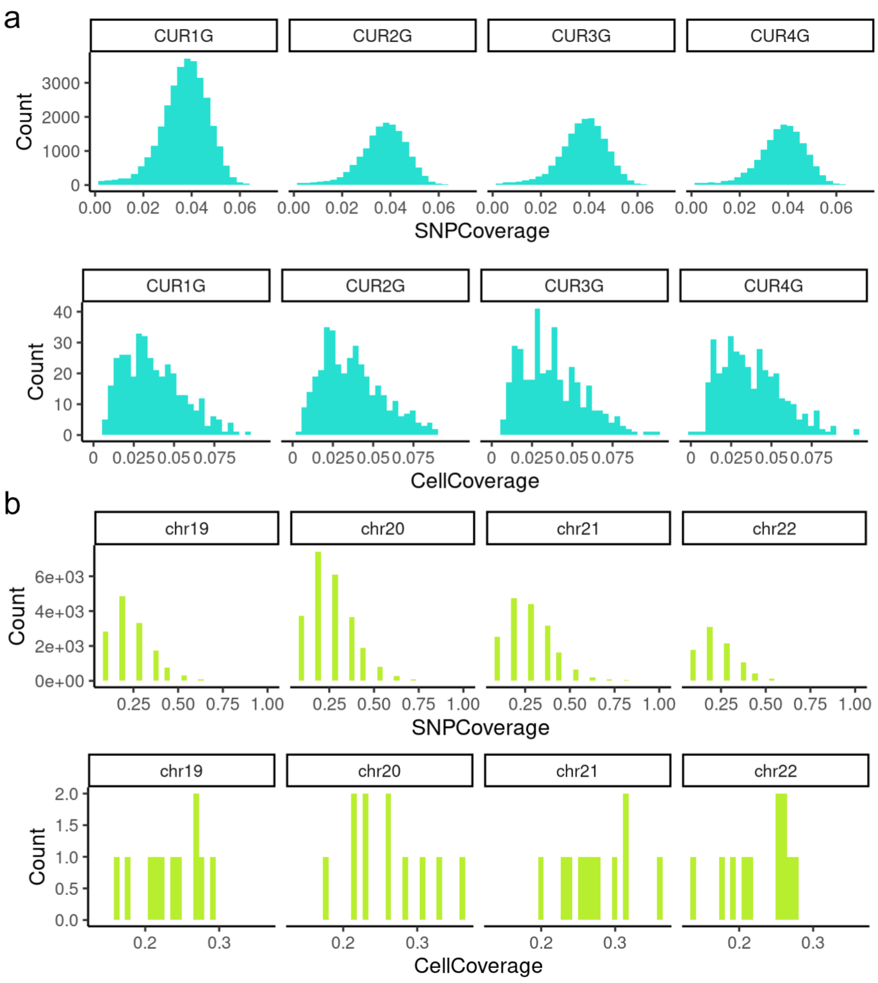

Figure S5: Dataset coverage comparison of human sperm dataset with apricot gametes The SNP density in the human sperm dataset is much higher comparing to the 10X scCNV apricot gamete dataset. **a)** Distribution of SNPs' coverage per cell and cells' coverage per SNP are plotted for apricot gametes [9]. **b)** Distribution of SNPs' coverage per cell and cells' coverage for the four smallest chromosomes in human genome for the human sperm dataset[10]

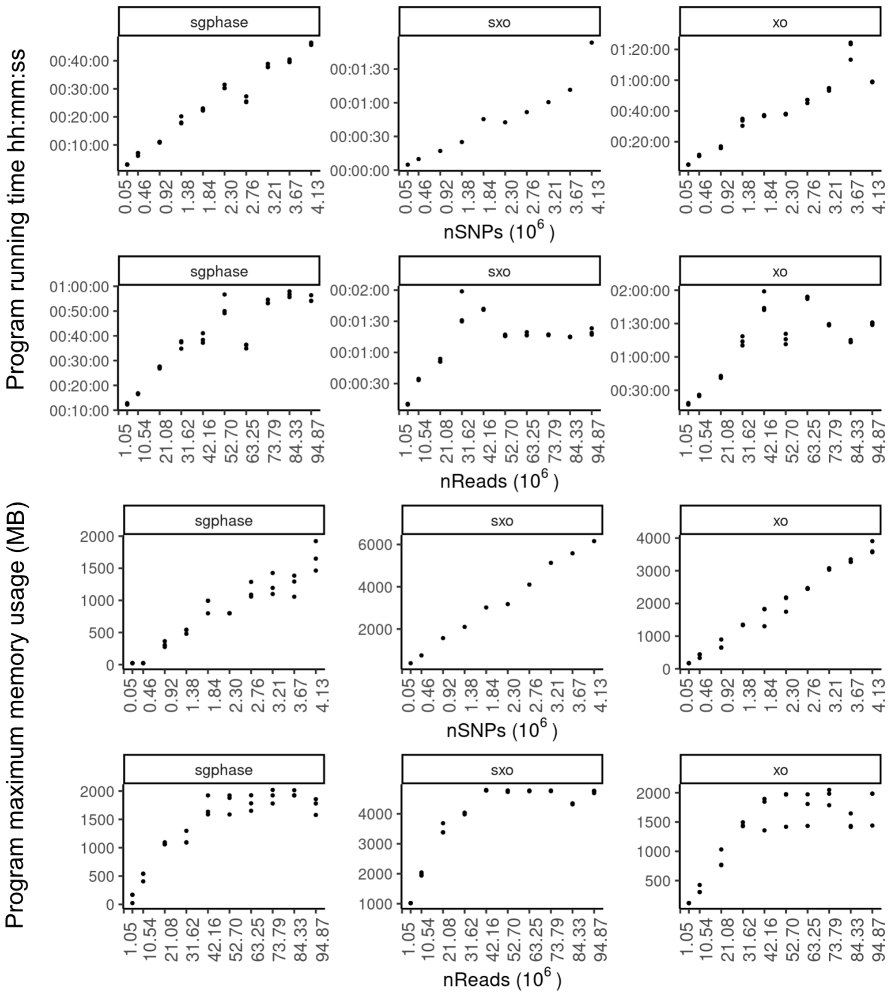

Figure S6: Module running time and memory usage by *sgcocaller* with varying input sizes . Top two panels show the running time in format of hour:minute:seconds of different modules with varying number of SNPs (nSNPs) and number of DNA reads (nReads) to process. X-axis labels are in the scale of $10^6$. Bottom two panels show the memory usage with the reported 'max_rss' in mega bytes (MB), which is maximum amount of memory occupied by a process at any time that is held in main memory (RAM), for different modules. Each measurement has been repeated three times by using the 'benchmark' function from `Snakemake` [11].
